# Stabilization of F-actin by *Salmonella* effector SipA resembles the structural effects of inorganic phosphate and phalloidin

**DOI:** 10.1101/2023.12.26.573373

**Authors:** Ewa Niedzialkowska, Lucas A. Runyan, Elena Kudryashova, Edward H. Egelman, Dmitri S. Kudryashov

## Abstract

Entry of *Salmonella* into host enterocytes strictly relies on its pathogenicity island 1 effector SipA. We found that SipA binds to F-actin in a unique mode in a 1:2 stoichiometry with picomolar affinity. A cryo-EM reconstruction revealed that SipA’s globular core binds at the grove between actin strands, whereas the extended C-terminal arm penetrates deeply into the inter-strand space, stabilizing F-actin from within. The unusually strong binding of SipA is achieved via a combination of fast association via the core and very slow dissociation dictated by the arm. Similarly to P_i_, BeF_3_, and phalloidin, SipA potently inhibited actin depolymerization by ADF/cofilin, which correlated with the increased filament stiffness, supporting the hypothesis that F-actin’s mechanical properties contribute to the recognition of its nucleotide state by protein partners. The remarkably strong binding to F-actin maximizes the toxin’s effects at the injection site while minimizing global influence on the cytoskeleton and preventing pathogen detection by the host cell.

## Introduction

Pathogenic *Enterobacteriaceae* are the leading cause of enteric and diarrheal diseases. The World Health Organization (WHO) estimates that *Salmonella* spp., *Shigella* spp., and *Escherichia coli* cause ∼380-1,360 million enteric illnesses and ∼350,000-920,000 deaths yearly worldwide^1^. Moreover, the emergence of multi-antibiotic resistance among these *Enterobacteriaceae* is a pressing concern in healthcare settings throughout the world, urging the development of new treatments to combat these pathogens. Understanding how *Enterobacteriaceae* invade host cells opens new targets for designing antimicrobial drugs.

SipA is a 74-kDa protein from *Salmonella* spp. that is encoded by *Salmonella* pathogenicity island 1 (SPI-1) and delivered to the host cell by type III secretion system 1 (T3SS1)^2^. Once recognized as a protein with a secondary role in *Salmonella* invasion whose function is to retain the bacterium at the host cell^3,4^, SipA has been recently reinstated as the most critical factor for invasion into polar epithelial cells under physiological conditions^5^. In contrast to the actin ruffle-dependent invasion of the pathogen into non-polar cultured cells that heavily depends on the injection of SopB, SopE and SopE2 effectors^3^, entry into polarized epithelial cells relies on SipA-dependent localized actin cytoskeleton rearrangements near the entry site^5^. The detailed role of SipA in this process and its ability to manipulate actin dynamics remains obscure.

SipA contains two well-defined globular α-helical N- (residues 48-264) and C-terminal (residues 512-658) domains flanked by disordered regions of various lengths^6,7^ (Fig. 1a). SipA’s N-terminal domain contains a bacterial chaperone InvB-binding region (residues 27-264) that facilitates translocation^7^ through the T3SS1 apparatus. This region encompasses the foci-forming F1 domain (residues 170-271)^8^, which contains the <u>s</u>oluble <u>N</u>-ethylmaleimide-sensitive factor <u>a</u>ttachment protein <u>re</u>ceptor (SNARE)-like peptide (residues 180-232) involved in remodeling of early endosomes^9^. This functional domain is followed by the sequence that stimulates transepithelial migration of polymorphonuclear leukocytes (residues 294-424)^10^, substantially overlapping with the SipA self-association domain (residues 280-394)^8^. SipA has been shown to activate caspase 3, which cleaves the toxin after Asp435 into N- and C-terminal fragments (SipA_N_ and SipA_C_, respectively)^11,12^. SipA_C_ interacts with F-actin^6^ and contains a globular core domain (residues 512-658) flanked by flexible N- and C-terminal regions, hereafter designated as Arm1 and Arm2, respectively (Fig. 1a). F-actin saturated with SipA has a lower critical concentration and is resistant to severing and depolymerization by actin depolymerizing factor (ADF)/cofilin^13,14^.

**Fig. 1.**
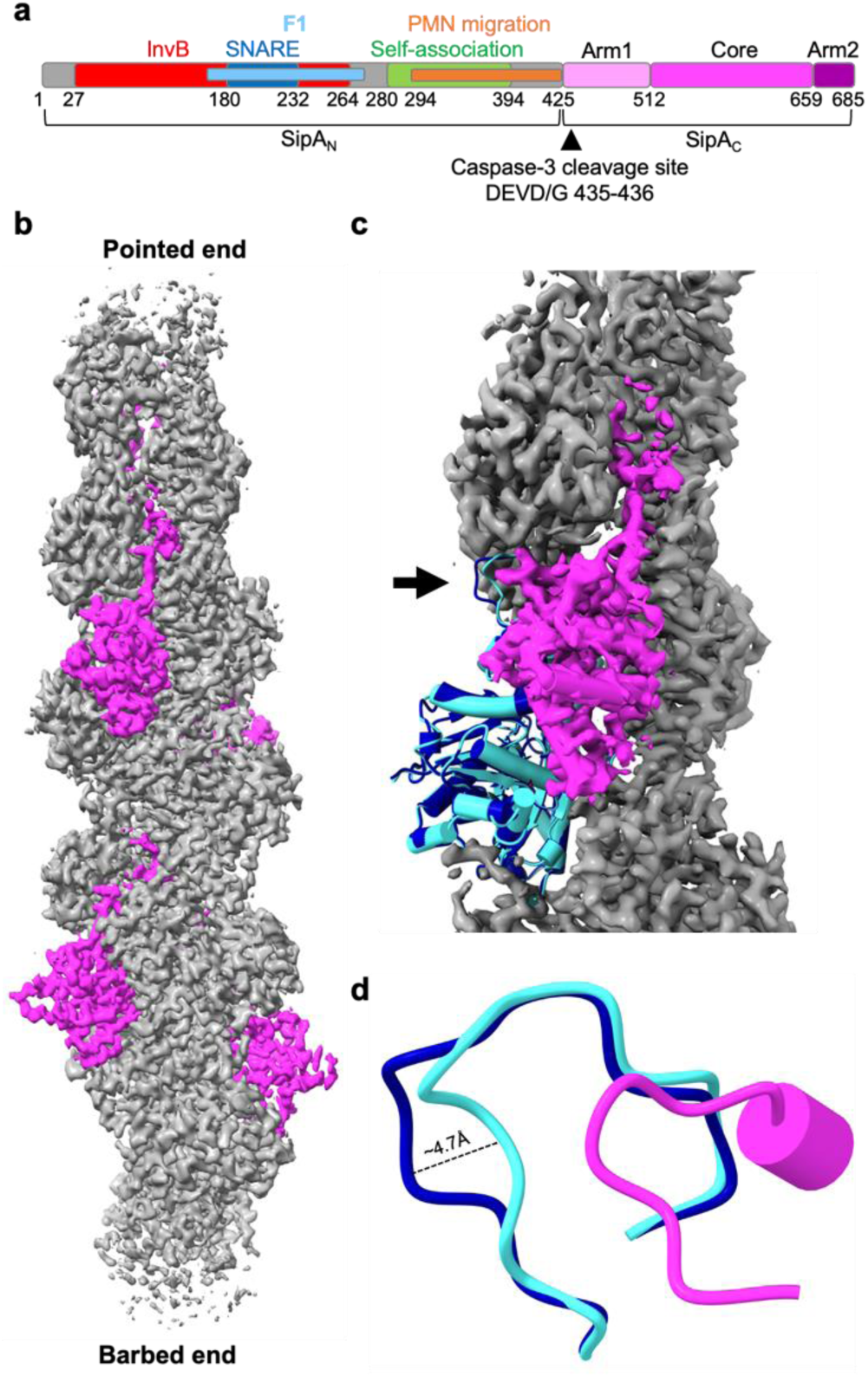
Cryo-EM reconstruction revealed nearest-neighbor exclusion of SipA_C_ on F-actin and minor perturbations to actin structure. **a,** SipA domain schematic; numbers represent amino acids in the SipA sequence (UniProtID: P0CL52). *InvB,* bacterial chaperone InvB-binding region (a.a. 27-264); *F1,* foci-forming domain (a.a. 170-271); *SNARE,* SNARE-like peptide (a.a. 180-232); *PMN migration,* stimulates transepithelial migration of polymorphonuclear leukocytes (a.a. 294-424); *Self-association,* SipA self-association domain (a.a. 280-394); *Core,* globular core domain (a.a. 512-658); *Arm 1* and *Arm2,* disordered regions flanking the Core (a.a 425-511 and 660-685, respectively). **b,** Cryo-EM density map of an actin filament (in grey) decorated by SipA_C_ (in magenta) (EMD-42161). **c,** Superposition of bare actin (blue; PDBID: 8D14) with SipA_C_/F-actin complex (SipA_C_ in magenta, actin monomer in cyan, F-actin density in grey) highlights changes in D-loop conformation (black arrow). **d,** Zoomed-in view of the D-loop region: the D-loop of protomer *n+1* (cyan) is displaced 4.7 Å upon SipA_C_ binding, relative to bare actin (blue). Displacement is measured between Cα atoms of actin Gly48.

The actin cytoskeleton is a highly dynamic system that mediates various cellular processes, such as cell motility, adhesion, intracellular transport, membrane remodeling, and contractile ring closure during cell division^15^. Monomeric actin (G-actin) polymerizes into filaments (F-actin)^16^, which can be further assembled into higher-order structures with numerous functionalities^17^. The balance between G- and F-actin is highly regulated^18^ and perturbations in G-/F-actin dynamics can be lethal. For example, cyclic peptides phalloidin and jasplakinolide produced by *Amanita phalloides* and *Jaspis spp.,* respectively^19,20^, manifest their toxicity by stabilizing F-actin. The key factors that mediate actin recycling are ADF/cofilin isoforms, which induce conformational changes in F-actin, promote its severing and, with the assistance of actin-interacting protein 1 (AIP1) and cyclase associated proteins 1 and 2 (CAP1/2), accelerate F-actin depolymerization^21-23^.

The interaction between the SipA^446-684^ fragment and F-actin was previously studied using negative-stain electron microscopy (EM), and stapling of the two actin strands with the fragment’s N- and C-terminal nonglobular arms was postulated^6^. It was suggested that such binding would lead to the increased mechanical stability of the more connected actin strands^6,24^, but the low resolution of the structure prevented revealing the atomic details of this interaction. To better comprehend the structural mechanisms of actin stabilization by SipA that is critical at early (invasion) and late (maintenance and protection of *Salmonella*-containing vacuole (SCV)) stages, we analyzed SipA-decorated F-actin using cryo-EM reconstruction and performed detailed biochemical characterization of the SipA/F-actin interaction, particularly at low-molar ratios of SipA to actin relevant for an infected cell.

## Results

### Cryo-EM revealed a 1:2 stoichiometry and bimodal binding of SipA to F-actin

Our initial attempts to reconstruct F-actin decorated by SipA^425-685^ (hereafter, SipA_C_) assuming a 1:1 ratio to actin were complicated by lower density for the toxin compared to actin, which translated into less than 50% SipA_C_ occupancy. Visual inspection of the map revealed continuous density arising from SipA_C_ that was connected between adjacent binding sites, which must be artifactual. At a 1:1 stoichiometry, the manually built C-terminal tail of SipA (Arm2) generated steric clashes with the globular domain of the next SipA_C_ in the adjacent binding site, prompting us to redo the reconstruction assuming a 1:2 ratio. A 3D variability analysis (3DVA) algorithm^25^ was applied to 2D classified particles followed by a round of 3D classification. Particles from a 3D class with no steric clashes between SipA molecules were refined with a 112.075 Å rise and −306.344° rise, corresponding to an asymmetric unit containing four actin subunits and two SipA_C_ molecules. Ultimately, non-uniform refinement was performed on 1,548,993 particles after the position and orientation of volumes and particles were adjusted with helical parameters 112.075 Å rise and −306.344° twist (Supplementary Fig. S1). The resulting 3.2 Å map (Fig. 1b, Supplementary Fig. S2a,b) contained well-defined density for almost all of actin and most of SipA_C_ with the resolution decreasing towards the periphery (Supplementary Fig. S2c,d). The C-terminal Arm2 could now be easily traced in the density extending from the globular domain without steric clashes with other SipA_C_ subunits.

The SipA_C_-decorated filament closely resembles the naked ADP-P_i_ actin (PDBID: 8D14^26^; RMSD 0.4 Å), and we were able to model ADP, inorganic phosphate (P_i_), and Mg^2+^ in the nucleotide-binding cleft (Supplementary Fig. S3a). The only difference was in the position of the DNaseI-binding loop (D-loop; Fig. 1c,d; Supplementary Fig. S3c), a highly dynamic, nucleotide-sensing element involved in establishing short- and long-pitch contacts in F-actin and in binding to several actin-binding proteins^27^. Binding of SipA_C_ caused a ∼5-Å displacement of Gly48 in the D-loop compared to a naked reconstruction of ADP-P_i_ actin (PDBID: 8D14; Fig. 1d).

The SipA_C_ construct used for cryo-EM reconstruction comprises an N-terminal unstructured arm (residues 425-511; Arm1), a globular region (residues 512-658; Core) composed of five helices linked by short loops, and a 26-residue C-terminal arm (residues 659-685; Arm2). Arm1 was not seen in the density, presumably due to disorder. Four of the Arm2 amino acids (residues 669 – 672) were only partially visible in the reconstruction. The globular domain of SipA_C_ bridges actin subunits both along the same strand and across the opposite strands. Counting from the barbed end of the actin filament, the SipA_C_-Core is nested between subdomain 4 of protomer n, subdomains 1 and 2 of *n*+1, subdomains 3 and 4 of *n*+2, and subdomain 1 of *n+3* (Fig. 2a). The actin-binding interface in the Core is formed by ionic interactions between actin Asp80 of protomer *n*+1 and Arg551 of SipA_C_ (Fig. 2b). Actin Glu224 and Glu316 of protomer *n*+2 form salt bridges with Arg628 and Lys618 of SipA_C_ (Fig. 2c,d). The SipA_C_/F-actin interface is also stabilized by hydrophobic interactions between Phe266 of actin *n+2* and Val532 of SipA_C_ (Fig. 2e) and intimate connections between Met538 and His539 of SipA_C_ and the D-loop of actin *n+1* (Fig. 2f). Met538 also contributes the side chain to a hydrophobic contact with actin Met355 of protomer *n*+3 (Fig. 2f), enforcing the longitudinal interprotomer contacts in F-actin. Among other D-loop contacts, the side chain of actin Gln41 is within a hydrogen bond distance from the carbonyl group of SipA’s His536, while the carbonyl of Val45 and side chain of Gln49 may form hydrogen bonds with His539 of SipA, assuming the latter is protonated (Fig. 2f).

**Fig. 2.**
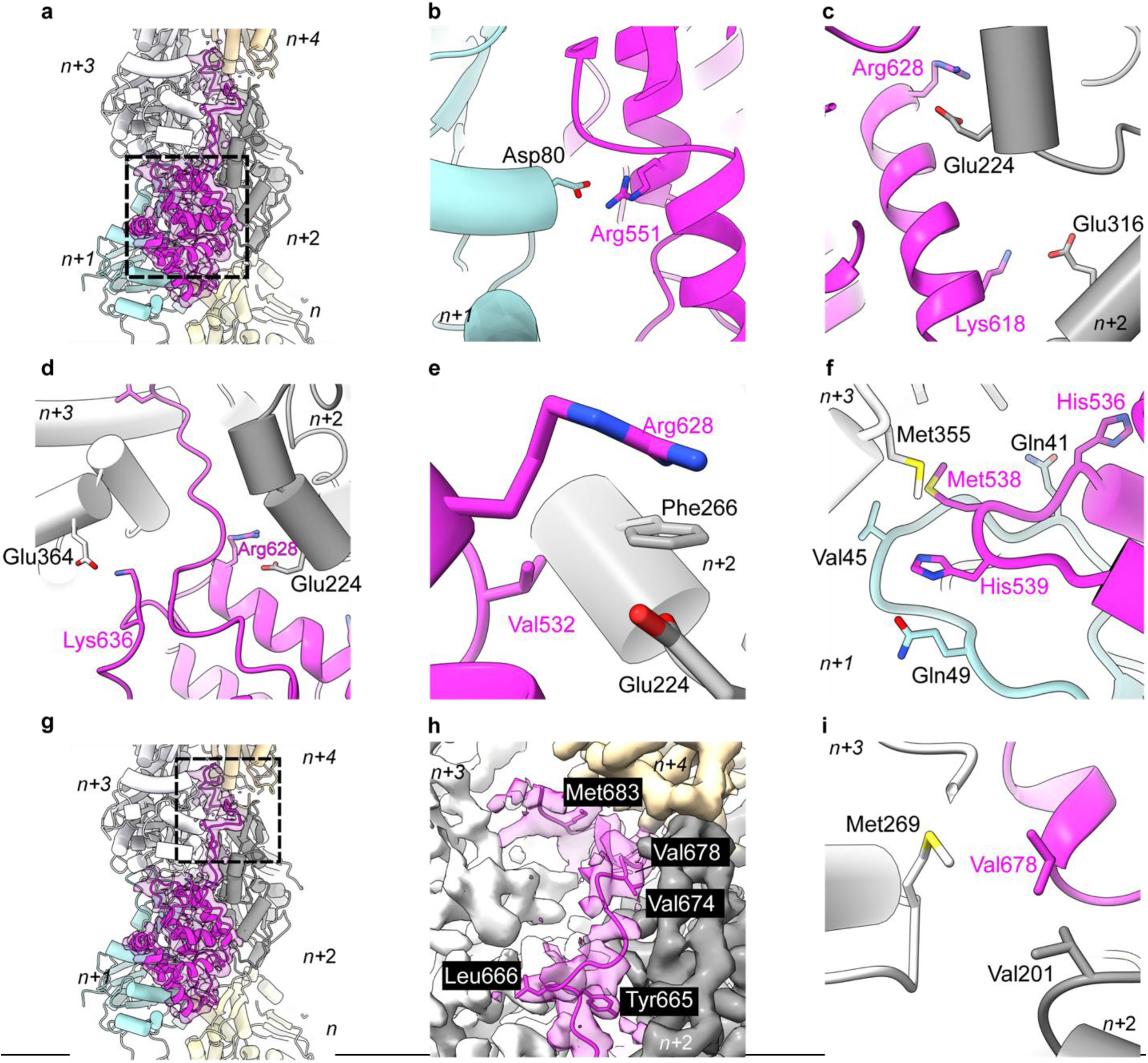
SipA_C_ binds actin via two distinct regions, the Core and Arm2. **a,** Atomic model fit to SipA_C_- decorated-actin cryo-EM density (EMD-42161). SipA_C_ (ribbon representation, magenta) contacts five actin protomers (cylinder representation, colored by protomer, numbered from the barbed end from *n* to *n+4*). Dashed box indicates SipA_C_-Core, whose interaction with actin is detailed in panels **b-f**. **b,** Actin protomer *n+1* residue Asp80 forms ionic bonds with Arg551 on SipA_C_. **c,** Actin protomer *n+2* residues Glu224 and Glu316 form ionic interactions with Arg628 and Lys618 on SipA_C_. **d,** Actin protomers *n+3* and *n+2* contribute ionic interactions from Glu364 and Glu224 to SipA_C_ Lys636 and Arg628 respectively, orienting the placement of Arm2. **e,** Phe226 on actin *n+2* is involved in hydrophobic interaction with Val532 of SipA_C_. **f,** Met538 of SipA_C_ interacts hydrophobically with Met355 on *n+3* protomer (white) while His539 H-bonds with carbonyl group of Val45 and Gln49 on *n+1* protomer (cyan) in the D-loop. Gln41 forms H-bond with His536. **g,** Overview of SipA/F-actin complex as in **a**, with the dashed box denoting Arm2-actin interactions (detailed in panels **h-i**). **h,** Hydrophobic residues Tyr665, Leu666, Val674, and Met683 on SipA_C_ anchor Arm2 to filament inter-strand space. **i,** Val678 on SipA_C_ contributes to inter-strand interactions by interacting with Val201 on protomer *n+2* and Met269 on protomer *n+3*.

### The Arm2 of SipA_C_ binds deep in the inter-strand cleft of F-actin

In our reconstruction, Arm2 is visible as an additional density deep between the two actin strands that cannot be explained by actin protomers or the SipA_C_-Core (Fig. 2g,h). Arm2 extends between the two actin strands and interacts with protomers *n+2*, *n+3* and *n+4*. The positioning of Arm2 into the inter-strand space is guided by ionic interactions between Arg628 of SipA_C_ and Glu224 of actin *n+2*, and Lys636 of SipA_C_ and Glu364 of actin *n+3* (Fig. 2d). Another residue crucial for proper docking of Arm2 is Tyr665, whose side chain points toward the helical axis of the actin filament, anchoring Arm2 between actin protomer *n+2* and *n+3*. Binding of Arm2 is further stabilized by hydrophobic contributions from Leu666, Val674, Val678 and Met683 of SipA_C_ (Fig. 2h). Val678 of SipA_C_ appears to be crucial for enforcing inter-strand interactions as it forms hydrophobic interactions with both Met269 of actin protomer *n+3* and Val201 of actin protomer *n+2* (Fig. 2i).

Almost all the residues involved in the interaction with SipA_C_ are conserved between skeletal actin, used in our reconstruction, and the non-muscle actin isoforms. The only exception is Val201 of skeletal actin, which is replaced by Thr201 in cytoplasmic actin. This substitution preserves the environment in the space between actin strands and should not affect SipA_C_ binding (Supplementary Fig. S3b).

### SipA_C_ differentially competes with phalloidin and jasplakinolide

Since our structural data contradicted a previously reported 1:1 stoichiometry of SipA_C_ to actin^13^, we independently confirmed the 1:2 molar stoichiometry in co-pelleting assays (Fig. 3a,b). Although the binding mode of SipA_C_ is unique for actin-binding proteins, the binding site of Arm2 within the inter-strand cleft overlaps with those of F-actin stabilizing toxins phalloidin and jasplakinolide (Fig. 3c,d)^20^. This observation prompted us to test whether phalloidin and jasplakinolide compete with SipA_C_ for F-actin. First, we assessed binding of both drugs to actin, using tetramethylrhodamine-isothiocyanate (TRITC)-phalloidin and SiR-actin, the fluorescent derivatives of phalloidin and jasplakinolide, respectively. We found the dissociation constant (K_d_) of TRITC-phalloidin binding to F-actin to be 45 nM (Fig. 3e) and that of SiR-actin to be 20 nM (Fig. 3f). Next, in competition experiments with SiR-actin, both SipA_C_ and SipA_C_-Core were able to fully displace jasplakinolide from actin, suggesting the competition of the drug with both Core and Arm2 (Fig. 3g,h). SipA_C_ construct competed notably more effectively, with K_d_ values derived from the binding isotherm competition equation being 0.18 nM and 81 nM, for SipA_C_ and SipA_C_-Core respectively (Fig. 3g). In competition with TRITC-phalloidin, SipA_C_-Core failed to displace the drug from actin (Fig. 3i), while SipA_C_ displaced only about half of the drug (Fig. 3j) with an apparent K_d_ of 5 nM (Fig. 3i), in agreement with *i)* 1:2 stoichiometry and *ii)* only the Arm2 and not the globular Core domain competing with phalloidin.

**Fig. 3.**
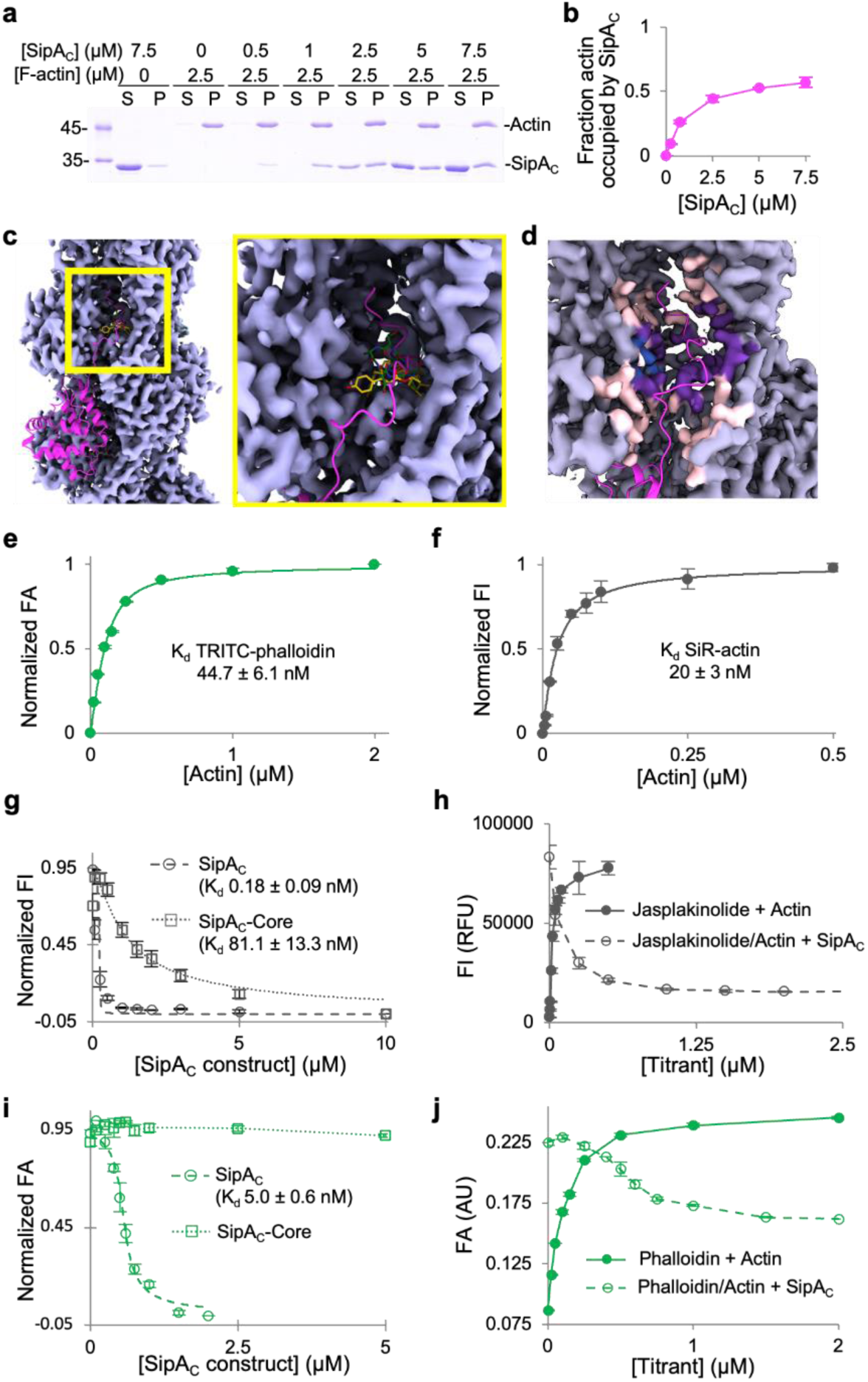
Arm2’s binding site on F-actin overlaps with those of phalloidin and jasplakinolide. **a,b,** Binding stoichiometry of SipA_C_ to F-actin was determined in co-sedimentation experiments. **a,** A representative SDS-PAGE co-sedimentation gel, where S and P designate supernatant and pellet fractions. **b,** Relative molar content of SipA_C_ to actin in the pellet fractions plotted as mean values ± SD from three technical replicates. **c,** An overview (left) and zoomed-in view (right) of the inter-strand space in the vicinity of SipA_C_-Arm2-binding site, showing overlap between Arm2 (magenta), phalloidin (green; PDBID: 6T1Y) and jasplakinolide (yellow; PDBID: 6T24). **d,** Light grey indicates density around actin residues that do not participate in SipA binding; light pink – density around residues that interact with SipA, but not with phalloidin or jasplakinolide; purple – density around residues that interact with both SipA and phalloidin/jasplakinolide; blue – density around residues that interact with phalloidin or jasplakinolide, but not with SipA. The C-terminal tail of SipA is in magenta. **e,** FA analysis of TRITC-phalloidin binding to actin. **f,** FI analysis of SiR-actin binding to actin. **g,** Normalized FI competition data for SiR-actin with SipA_C_ or SipA_C_-Core. **h,** Raw FI data for SiR-actin binding to F-actin and competition with unlabeled SipA_C_. **i,** Normalized FA competition data for TRITC-phalloidin with SipA_C_ or SipA_C_-Core. **j,** Raw FA data for TRITC-phalloidin binding to actin and competition with unlabeled SipA_C_ show that only ∼50% of TRITC-phalloidin is displaced by SipA_C_.

### Equilibrium binding revealed distinct roles of SipA_C_ domains in binding to actin

To reveal the roles of Core, Arm1, and Arm2 in SipA_C_ binding to actin, we conducted direct and competition fluorescence anisotropy measurements with fluorescein maleimide (FM)-labeled SipA_C_, SipA_C_-ΔArm2 and SipA_C_-Core constructs (Supplementary Fig. 4a). Identical, except unlabeled, constructs were used in the competition experiments. The presence of Arm2 notably increased the affinity of the toxin to actin (*e.g.,* compare ∼0.9 nM and 120 nM K_d_s for the labeled SipA_C_ and SipA_C_-ΔArm2, respectively; Supplementary Table S1 and Supplementary Fig. 4b-e). In contrast, the presence of Arm1, which was disordered in the cryo-EM reconstruction, decreased the binding strength of SipA_C_ by a factor of ∼4-10, depending on the experimental approach (Supplementary Table S1, Supplementary Fig. 4d-g). This inhibition is likely due to the entropic cost of Arm1’s restricted flexibility in the actin-bound state. Competition experiments overall confirmed these findings while showing even higher affinities of the unlabeled constructs for actin (Supplementary Fig. 4e,g). Measuring these high affinities necessitated dropping actin concentrations below its critical concentration for polymerization (C_C_ ∼100 nM). To overcome this challenge, the ΔArm2 constructs (*i.e.*, SipA_C_-Core and SipA_C_-ΔArm2, which do not compete with phalloidin) were titrated with phalloidin-stabilized actin, while the SipA_C_ construct (competes with phalloidin but also potently stabilizes F-actin) was titrated by free F-actin upon its rapid dilution from concentrations above critical. Since such limitations and variations are vulnerable to inaccuracies, and to get further insight into the contribution of SipA_C_ domains to F-actin binding, we reassessed the binding by measuring its kinetic characteristics.

### Kinetic experiments suggest a bimodal “docking-then-anchoring” binding of SipA_C_ to actin

We monitored changes in anisotropy of FM-labeled SipA_C_ constructs upon their binding to or competitive dissociation from F-actin using a stopped-flow fluorometer (Fig. 4). Each SipA_C_ construct had a linear dependence of *k_obs_* on F-actin, yielding *k_on_* values within a ∼3-fold range of each other (Fig. 4a, Supplementary Table S1). SipA_C_-ΔArm2 had the lowest association rate of 1.4 µM^-1^s^-1^, which increased to 3.7 µM^-1^s^-1^ upon removal of Arm1. The presence of Arm2 accelerated binding to 3.8 µM^-1^s^-1^ even in the presence of Arm1, thereby compensating for the binding deficiency caused by Arm1. Interestingly, Arm2 alone, synthesized as a peptide with an N-terminal fluorescein isothiocyanate (FITC) label, failed to bind actin in anisotropy experiments. Additionally, the Arm2-only constructs showed diffuse localization upon expression in cells (Fig. 5) regardless of the fluorescent protein tag location (Fig. 5a,h,i). Taken together, these results demonstrate the critical role of the globular core domain in the initial docking of SipA to F-actin.

**Fig. 4.**
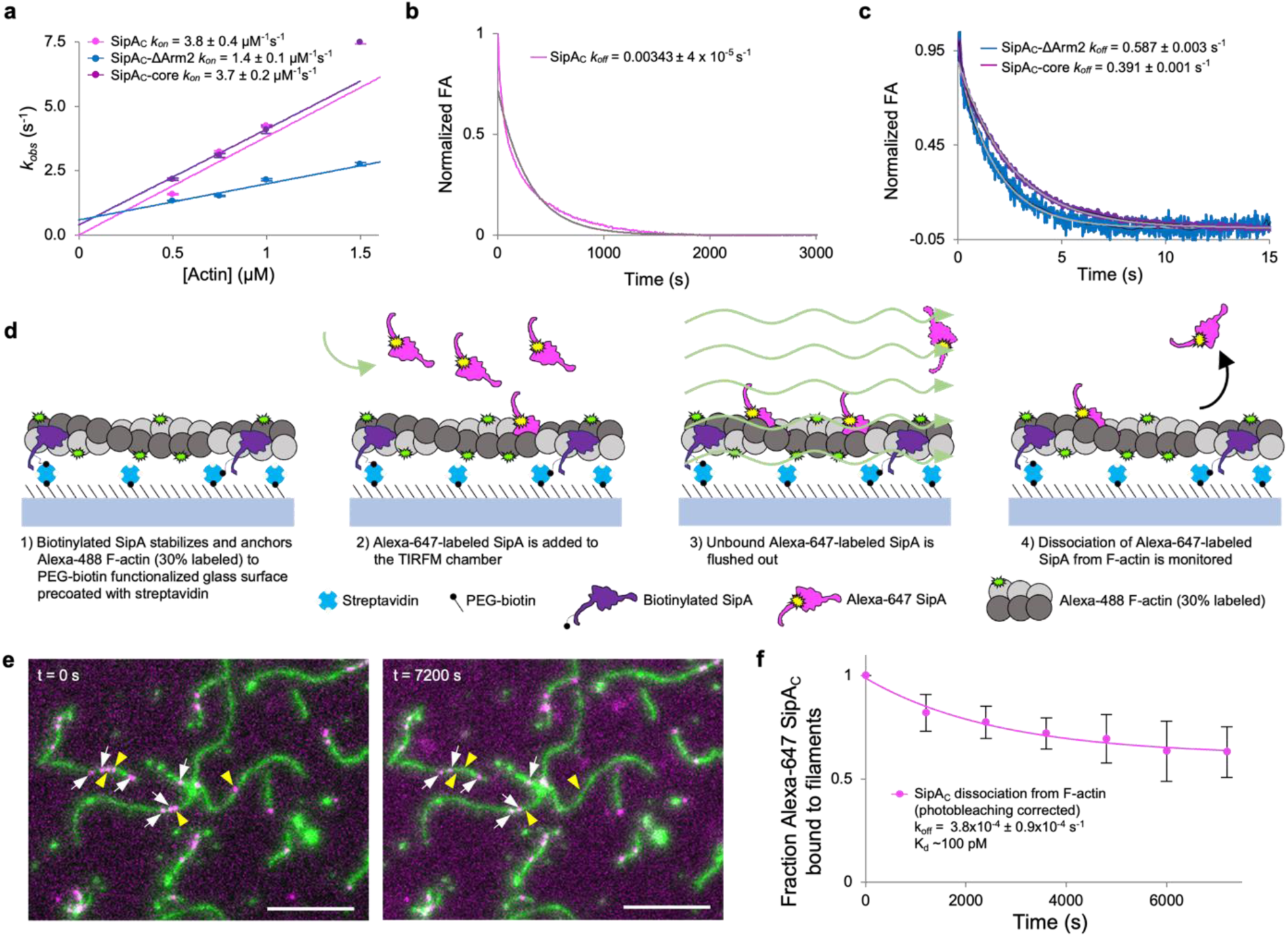
Association and dissociation rates of SipA_C_ are dictated by the Core and Arm2 domains, respectively. **a,** The *k_obs_* values determined from kinetic association experiments plotted against actin concentration show a linear relationship and yield *k_on_* values for each SipA_C_ construct. **(b,c)** Dissociation kinetics of FM-SipA_C_ (**b**), FM-SipA_C_-ΔArm2 and SipA_C_-Core (**c**) from F-actin upon competition with excess of respective unlabeled constructs yields *k_off_* values for each construct. Grey traces represent fits to a single exponential. (**d**) Schematic of TIRFM experiments to monitor dissociation of SipA_C_ from F-actin. (**e**) Representative images from TIRFM dissociation experiments. White arrows indicate Alexa-647 SipA_C_ molecules present throughout the experiment duration from t = 0 s (left) to t = 7200 s (right); yellow arrowheads indicate Alexa-647 SipA_C_ molecules present at t = 0 s but not at t = 7200 s. (**f**) Dissociation of Alexa-647 labeled SipA_C_ from Alexa-488 labeled F-actin in TIRFM corrected for photobleaching was fit to a single exponential decay equation providing an estimate of k_off_, which yields a K_d_ of ∼100 pM.

**Fig. 5.**
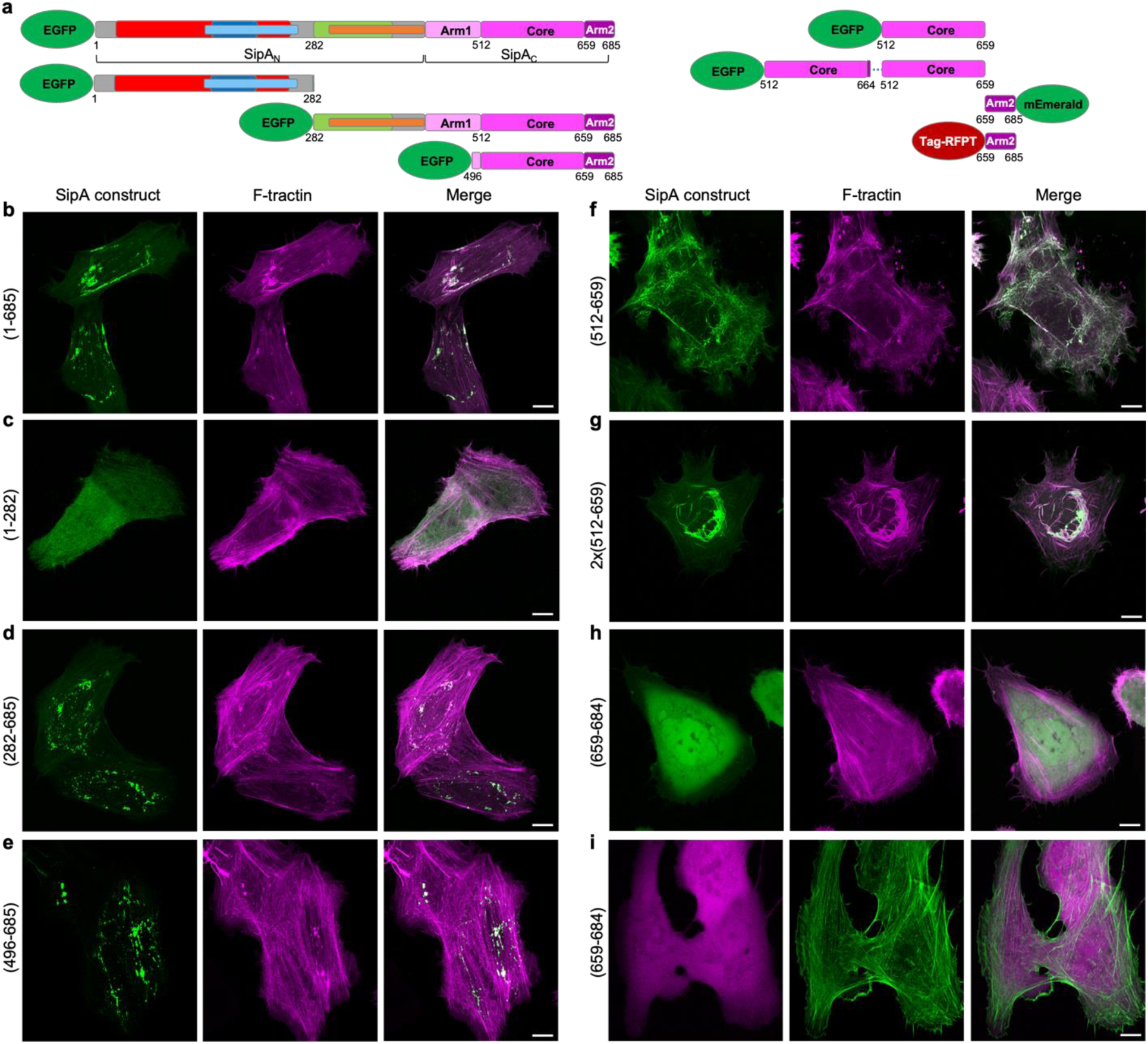
Intracellular localization of SipA constructs. **a,** Domain schematics of SipA constructs used for transfections. **b-i,** Representative maximum intensity projection images of Z-stacks of U2OS cells co-transfected with the SipA constructs and F-tractin (to visualize actin). Scale bars are 10 µm.

In a classical experimental design described above, the dissociation rate constants (*k_off_*) was determined by adding an unlabeled competitor to the complex of the labeled molecule (*e.g.,* FM-SipA_C_) and its partner (*e.g.,* actin). The competitor must be added in a sufficient, empirically determined excess to prevent reassociation of the studied molecule after its spontaneous dissociation. While this method provides reliable outcomes for simple, single-domain interactions (*e.g.,* SipA_C_-core with actin), it typically reports overestimated rates for more complex situations when two or more domains are involved^28^. With these limitations in mind, we determined that the constructs missing Arm2 showed similar dissociation rate constants (*k_off_*) of 0.39 and 0.59 s^-1^ whereas SipA_C_ displayed a 150-fold slower rate of 3.4×10^-3^ s^-1^(Fig. 4b,c). Yet, even this slow rate is likely to be overestimated as the competition with the unlabeled protein erases the key property of dual valency/avidity interactions. Indeed, in avidity interactions involving two or more binding sites, the dissociation of one of the domains is more likely to be followed by its reassociation than by the dissociation of the entire complex^28^. Instead, the excess of unlabeled SipA_C_ takes the place of the dissociated domain, preventing its rebinding and resulting in much faster dissociation rates overall.

To address this experimental deficiency, we directly measured the dissociation of SipA_C_ by counting the drop in actin-associated Alexa-647-SipA_C_ molecules by taking snapshots in 20-minute intervals for 2.5 hours in TIRF microscopy experiments (Fig. 4 d-c). The dissociation rate determined by this approach was 3.8×10^-4^ s^-1^, resulting in an equilibrium dissociation constant of 100 pM (Fig. 4 c). The *k_off_* of SipA_C_ is similar to that of phalloidin^29^, but the overall binding of SipA_C_ is ∼170 folds stronger mostly due to much slower association rate of phalloidin (0.028 µM^-1^s^-1^)^29^ as compared to SipA_C_ (3.8 µM^-1^s^-1^). This comparison reveals the importance of both the Core and Arm2 domains in attaining this unprecedented interaction strength by “docking” (via Core) and “anchoring” (via Arm2) SipA_C_ to actin. Whereas the former provides fast association, the latter ensures slow dissociation, resulting in the unique actin-binding properties of SipA_C_.

The detected fast binding and remarkably slow dissociation from actin explains the localization of SipA at the site of injection by *Salmonella*, as SipA’s diffusion inside the host cell is expected to be negligible under such conditions. Accordingly, all SipA constructs containing both the Core and Arm2 actin-binding domains were localized in actin-enriched compact foci (Fig. 5b,d,e), likely depicting the sites of their synthesis on ribosomes. Besides the sites of SipA localization, the geometry of actin fibers in such cells was not substantially affected. The constructs lacking both actin-binding domains or containing only Arm2 showed diffused localization (Fig. 5c,h,i). Interestingly, the SipA_C_-Core domain lacking Arm2 localized along actin fibers throughout the entire cell area, notably perturbing the geometry of the actin cytoskeleton (Fig. 5f). Finally, an artificial construct containing two tandem SipA_C_-Core domains induced profound bundles primarily in the perinuclear area of the cells (Fig. 5g). Therefore, the unique contribution of both Core and Arm2 ensures compact SipA localization and limits the generalization of its toxic effects within the cell.

### Arm2 enables cooperative stabilization of actin by SipA_C_

To evaluate how such strong binding may affect the dynamics of actin and its actin-binding partners, we explored the thermal stability of F-actin and its depolymerization/severing by ADF. It is known that the stoichiometric doses of SipA protect F-actin against depolymerization by cofilin^14^. Yet, given the abundance of actin in the cell, stoichiometric toxin concentrations cannot be considered physiological and are insufficient for understanding the pathogenesis^30-32^. We found that the SipA_C_ constructs containing Core but lacking Arm2 caused no stabilization of actin in thermal denaturation experiments (Fig. 6a). In contrast, SipA_C_ raised the melting temperature of F-actin by ∼6°C, as compared to ∼10°C stabilization by phalloidin (Fig. 6a). The observed stronger actin stabilization by phalloidin was likely due to two factors. First, 1:1 stoichiometry of phalloidin to actin enforces the inter-strand connections between all actin subunits. Second, a lower thermal stability of SipA_C_ may prevent actin stabilization at higher temperatures.

**Fig. 6.**
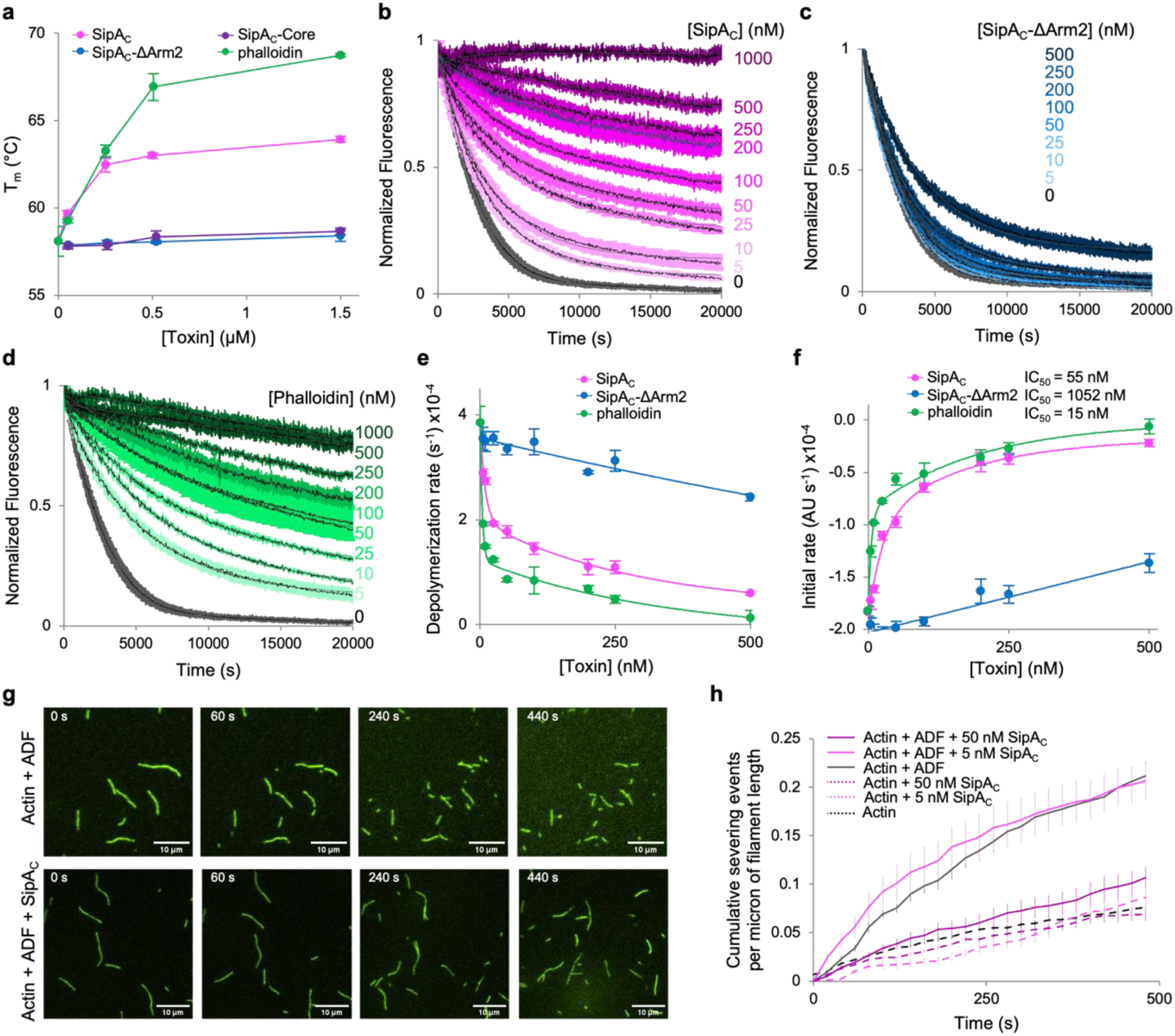
Arm2 of SipA_C_ mimics the effects of phalloidin on actin stabilization. **a,** Effects of bound SipA_C_ constructs or phalloidin on melting temperature (T_m_) of F-actin measured by differential scanning fluorimetry. **b-d,** Inhibition of ADF-stimulated actin depolymerization by various concentrations of SipA_C_ (**b**), SipA_C_-ΔArm2 (**c**), and phalloidin (**d**). **e,f,** Data shown in **b-d** were quantified as single-exponential decay rates (**e**) or initial depolymerization rates (**f**) and plotted against toxin concentration. Lines represent double exponential decay fits for SipA and phalloidin, and linear fits for SipA_C_-ΔArm2. **g,h,** F-actin severing analysis. **g,** Representative TIRF microscopy images of F-actin severed in the presence of ADF alone (top panels), or ADF and SipA_C_ (bottom panels). **h**, Quantification of severing events plotted over time.

In the cell, F-actin depolymerization is indispensable for proper actin dynamics and is mediated largely by ADF/cofilin proteins and their partners. In reconstituted ADF-mediated actin depolymerization experiments using pyrene-labeled actin (Fig. 6b-f), the protective effects of SipA_C_ and phalloidin were observed at the lowest tested (1:400) mole ratio to actin, while 50% inhibition (IC_50_) of depolymerization was achieved at 1:36 and 1:133 mole ratios, respectively (Fig. 6b,d,f). Like in the thermal denaturation experiments (Fig. 6a), the effects of phalloidin were similar but more pronounced, leading to overall slower depolymerization rates throughout the entire range of toxin concentrations. In both cases, the inhibition of depolymerization was clearly biphasic and could be fit well to dual exponential decay equations (Fig. 6e). Notably, the SipA_C_-ΔArm2 construct caused much less inhibition of depolymerization with a single slope reminiscent of that of the second component in the stabilization curves by phalloidin and SipA_C_ (Fig. 6e).

Since the accelerated depolymerization by ADF is a function of both the faster dissociation of subunits from the filament end, and a larger number of ends produced due to severing, we directly assessed severing of F-actin by ADF in total internal reflection fluorescence microscopy (TIRFM) experiments (Fig. 6g,h). To explore the severing of actin filaments homogeneously decorated by SipA_C_ at the desired levels of saturation, the construct was incubated with pre-polymerized aged F-actin prior to applying the mixture into the TIRFM chamber. At a 1:200 mole ratio to actin (5 nM SipA_C_, 1 µM actin), SipA_C_ caused no measurable protection against severing by ADF (Fig. 6h, Supplementary Fig. S5). However, when SipA_C_ was present at a 1:20 mole ratio to actin (50 nM SipA_C_, 1 µM actin), decoration of actin by SipA_C_ reduced actin severing by ADF to nearly background levels (Fig. 6g,h, Supplementary Fig. S5, and Supplementary Movie S1), pointing to a notable cooperativity of the protective effect.

### Arm2 of SipA_C_ cooperatively increases F-actin stiffness

Partial decoration of F-actin by SipA_C_ may inhibit depolymerization at the filament ends due to the “capping” effect when the filament is shortened down to the nearest toxin-bound actin subunit. The inhibition of severing by sub-stoichiometric doses of SipA_C_, however, is less expected as it requires the effects of SipA_C_ binding to propagate to undecorated actin subunits. In the absence of obvious structural differences between ADP-P_i_ vs ADP actin filaments, a difference in their stiffness was recently proposed to account for a striking difference in their affinities for ADF/cofilin^26^. To explore whether the observed cooperative inhibition of depolymerization by ADF may have a similar nature, we measured the stiffness (expressed as persistence length (L_p_)) of actin filaments in their different nucleotide states and upon interaction with SipA_C_. For the filaments stabilized during their polymerization (ADP-P_i_ enriched, young filaments), saturating concentrations of SipA_C_ raised the L_p_ from 14.6 to 22.5 µm, while SipA_C_-ΔArm2 only modestly increased L_p_ to 15.7 µm (Fig. 7a). Similarly, SipA_C_ increased L_p_ of pre-polymerized actin filaments (ADP-enriched, aged actin) from 12 to 17.7 µm, which is comparable to the effect of phalloidin (17.8 µm) and notably stronger than the stabilization by a P_i_-mimic beryllium trifluoride (BeF_3_; 13.8 µm) (Fig. 7b). Importantly, the concentration dependence of the rise in stiffness caused by SipA_C_ correlated with that of the protection against depolymerization by ADF (Fig. 7b,c). This finding supports the hypothesis that the nucleotide-dependent binding of ADF/cofilins to F-actin may be controlled by the difference in stiffness between the ADP-P_i_ and ADP actin states^26^.

**Fig. 7.**
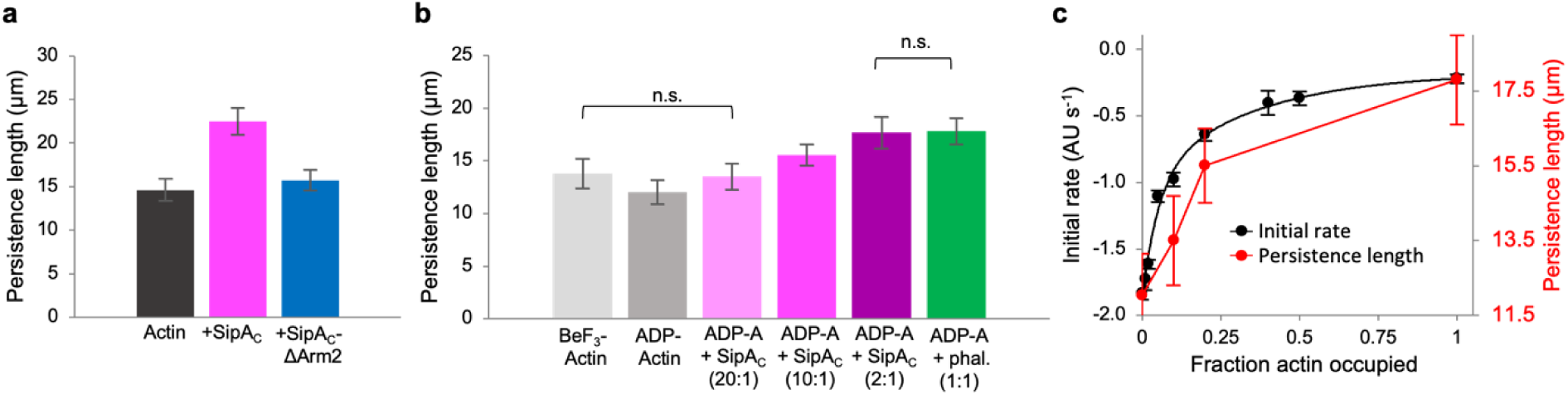
Actin filament stiffness is increased by SipA_C_ constructs containing Arm2. **a,b,** Persistence length (Lp) of actin filaments were measured as described in the Methods, and significant differences between average L_p_ values were determined by ANOVA analysis (Supplementary Table S2). **c,** Lp of ADP-actin (red) and the initial rates of depolymerization from Fig. 6f (black) were plotted as functions of actin filament saturation by SipA_C_, assuming the 2:1 binding stoichiometry.

### SipA_C_ moderately accelerates actin polymerization and strongly inhibits P_i_ release

Given the ability of a single SipA_C_ molecule to bridge up to five actin subunits, the toxin may stabilize actin nuclei during the nucleation process and thus promote polymerization. Also, the increased filament stiffness may affect the release of inorganic phosphate (P_i_) due to reduced thermal fluctuations. Since SipA_C_ binding to pyrene-labeled actin quenches its fluorescence, we opted to measure actin polymerization by light scattering. Both SipA_C_ and SipA_C_-ΔArm2 similarly promoted polymerization, suggesting that the stabilization of actin nuclei is dominated by fast binding of the Core (Fig. 8a). While both constructs also slowed the P_i_ release from polymerizing actin, the inhibition caused by SipA_C_ was notably stronger. Therefore, while the modest acceleration of polymerization is largely due to SipA_C_-Core, the inhibition of P_i_ release is dominated by the effects from Arm2. As the result of these two effects, actin filaments polymerized in the presence of SipA_C_ retain the ADP-P_i_ state longer than either naked filaments or even SipA_C_-ΔArm2-decorated filaments (Fig. 8b), which explains the well-defined density of P_i_ in our reconstruction of SipA_C_-decorated actin (Supplementary Fig. S3a).

**Fig. 8.**
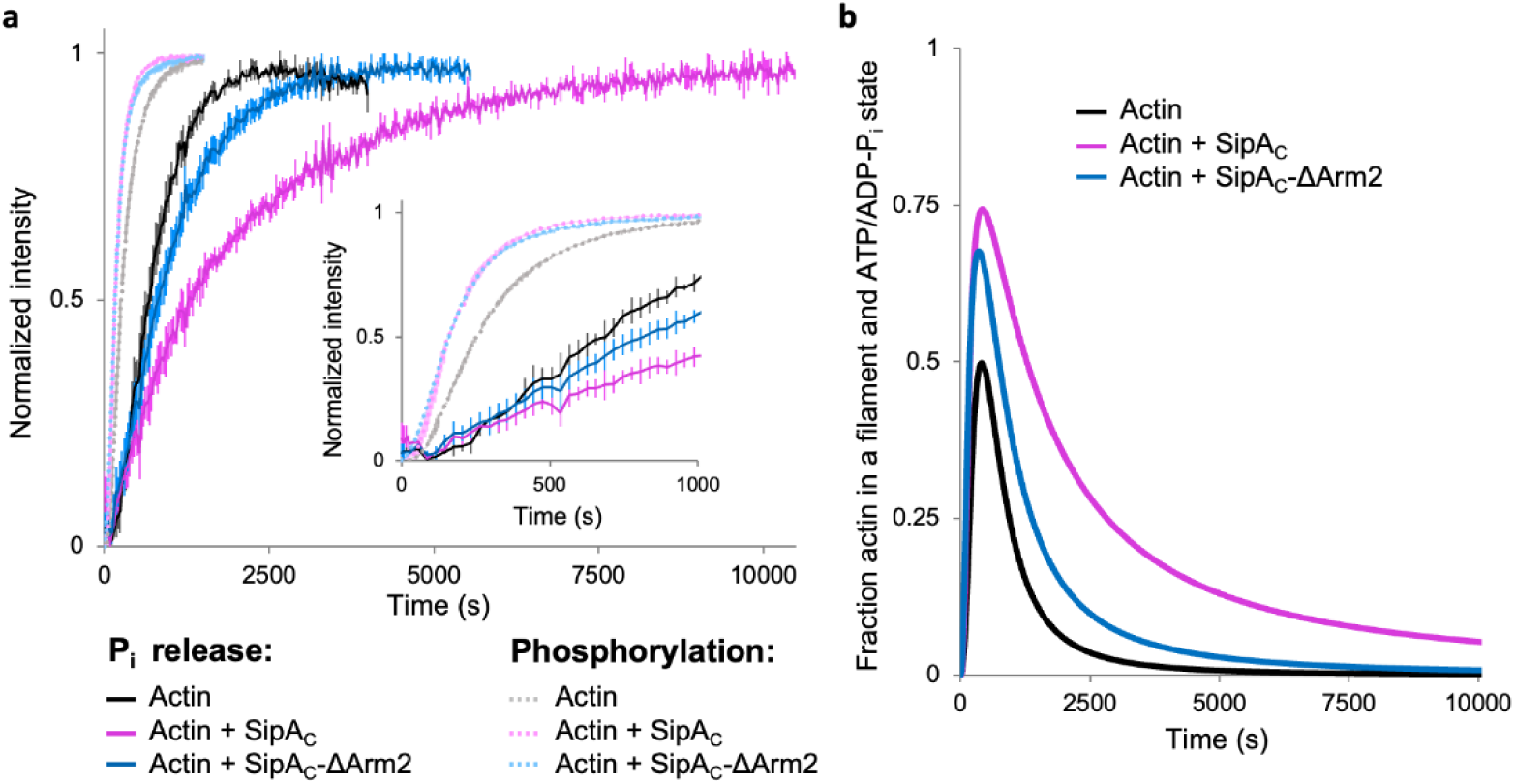
SipA_C_ accelerates actin polymerization and inhibits P_i_ release. **a,** Actin polymerization (light shade curves) and P_i_ release (dark shade curves) in the absence or presence of SipA_C_ and SipA_C_-ΔArm2. Inset emphasizes the early time points of the graphs. **b,** The fraction of filamentous actin in the ATP and ADP-P_i_ states was calculated by subtracting the normalized P_i_ release curves from actin polymerization curves (see Methods).

## Discussion

In this work, we discovered that binding of *Salmonella* effector SipA to F-actin represents a novel mode of actin interaction with its partners. Our atomic model based on the 3.2-Å resolution cryo-EM reconstruction indicates that SipA binds F-actin via extensive interactions with both the C-terminal globular Core domain and flexible Arm2. These contacts endow SipA_C_ with a picomolar K_d_ for F-actin, which is unprecedented for any known actin-binding partner. This interaction is granted by *i)* fast binding via the Core (”docking”) and *ii)* remarkably slow dissociation due to “anchoring” of SipA on actin via Arm2, penetrating deep into the inter-strand space. Strong binding of phalloidin (and likely jasplakinolide) is also achieved due to slow dissociation^33^, characteristic for the inter-strand binding mode shared by all three molecules^19,20^. Yet, a much faster association of SipA_C_, bestowed by its Core, results in ∼170-fold stronger binding, compared to the drugs. Therefore, SipA_C_ overcomes the limited accessibility of the inter-strand space^33^ by a priming interaction at the filament surface. Notably, Arm2 alone does not appreciably bind to actin (Fig. 5h,i), possibly due to a high entropic price for the association of a flexible peptide and a deeper location of the binding surface within the cleft (Fig. 3c,d).

An actin-binding domain of *Vibrio parahaemolyticus* VopV was also reported to compete with phalloidin and was found at the inter-strand space, with partial density localized within the cleft^34^. However, the low 9.6 Å resolution of the VopV-actin structure precluded accurate modeling of the peptide. The reported nanomolar K_d_ of the VopV repeat to actin is more like that of phalloidin and jasplakinolide, than SipA_C_. Whether this difference is due to a weaker external “docking” of VopV’s disordered domain or less deep interaction with the cleft remains to be established.

We confirmed a previous finding that SipA_C_ stabilizes F-actin against depolymerization by ADF/cofilin^14^. Moreover, we showed that this stabilization extends to physiologically relevant, low stoichiometric ratios of SipA to actin (Fig. 6), likely achievable upon actual infection. This stabilization is comparable to that caused by phalloidin, which, together with a much weaker effect of SipA_C_-ΔArm2, suggests that in both cases ADF/cofilins are inhibited by the peptides bound in the inter-strand space. This space is normally occupied by water molecules that provide discrete hydrophilic interactions, lubricate the contacts, and facilitate mechanical rearrangements required for filament flexibility^26^. Their replacement by strong inter-strand hydrophobic bridges provided by Arm2 or phalloidin explains the observed filament stiffening and stabilization effects *in vitro*.

The observed P_i_-release inhibition (Fig. 8a) can result from filament stiffening by Arm2 or by directly influencing the “back door” actin residues gating the P_i_ release (*i.e.,* methylated His73me, Gly74, Asn111, Arg177)^35^, or both. Indeed, Arm2 affects the His73me side chain orientation (Supplementary Fig. S6). The His73 of actin is subjected to posttranslational methylation by SETD3 methyltransferase^36^. Highly conserved among eukaryotes, N3-methylation of His73^36,37^ stabilizes the positive charge on histidine, which in turn enhances the dipole electrostatic interactions involving the carbonyl of G158, leading to strengthening of the hydrogen bond between carbonyl of G158 and side chain of His73me. As a result, the G158-V159 peptide bond is polarized, increasing the positive nature of the V159 backbone nitrogen. Such a chain of electrostatic interactions strengthens the V159-γ-phosphate hydrogen bond, that in turn may decrease the hydrolysis rate of ATP bound to F-actin^38,39^. In the SipA/F-actin complex, this chain of electrostatic interactions and hydrogen bond network around G158 and V159 are disrupted because SipA His682 causes the His73me side chain to rotate. Whether these changes affect ATP hydrolysis rates or P_i_ dissociation in the decorated filament remains to be found.

In the context of *Salmonella* infection, the remarkably strong binding to actin prevents diffusion of SipA away from the injection site, while also cooperatively stiffening partially decorated actin filaments, stabilizing them against depolymerization by ADF/cofilin. Combined, these effects enable highly effective local stabilization of the cortical actin by minute amounts of SipA, while also avoiding generalized effects on the actin cytoskeleton. The latter would be essential to prevent early detection of the pathogen by innate immune mechanisms, which can initiate inflammation and programmed cell death, leading to elimination of the pathogen. These speculations are supported by a normal morphology of the cells expressing SipA constructs containing both SipA_C_-Core and Arm2, which are localized in distinct foci (Fig. 5b,d,e), and notably stronger cytoskeletal disturbance in the cells expressing the SipA_C-_Core missing Arm2 (Fig. 5f,g).

Whereas tight binding of SipA_C_ to actin enables highly confined anchorage to the cytoskeleton, similarly effective connection with the bacterium is provided by the F1 domain encompassing residues 170-271, which alone is sufficient to form *Salmonella*-localized foci during the toxin injection^8^, but not upon expression in host cells (Fig. 5c). The mechanisms behind the formation of these foci remain unclear but likely involve membrane remodeling via association with integral membrane proteins, as an overlapping 180-232 a.a. peptide mimics a host SNARE peptide in establishing contacts with syntaxin-8 to promote fusion with early endosomes^9^. Both the actin-binding domain and F1 peptide of SipA are essential for the toxin-mediated *Salmonella* entry^8^.

SipA interaction with actin does not occlude the binding sites of major ABPs (tropomyosin, myosins, calponin-homology domain proteins, *etc.*), and thus SipA may involve host actin-binding proteins to support *Salmonella* invasion. However, host proteins whose binding requires major rearrangement of actin contacts, such as ADF/cofilins, or host proteins that also bind at the groove between actin strands such as the Arp2/3 complex and coronins will likely interfere with SipA binding. SipA has been shown to cooperate with plastin, a versatile host protein involved in crosslinking actin filaments in various types of actin bundles and assemblies^40^. SipA increases the bundling of actin by plastin and recruits the latter to the site of *Salmonella* invasion^41,42^. Superposition of F-actin structures decorated by SipA_C_ and by an actin-binding domain of plastin^40,43^ shows no steric conflicts between these proteins (Supplementary Fig. S7). Since no direct interaction between SipA and plastin has been detected^41^, the bundling potentiation is likely mediated by F-actin. The longer and stiffer (and, therefore, straighter) SipA-decorated filaments might be a better substrate for crosslinking by most, if not all, actin-bundling proteins. Finally, SipA may serve as a partner protein that weakens the inhibitory association between actin-binding domains of plastin^40^, favoring the bundling-competent open state^44^.

While SipA and, with the above reservations, VopV are the only proteins currently recognized to bind in the cleft between actin strands, a very high affinity and increased filament stability granted by such a mode are likely to be beneficial for specific cytoskeleton compartments of a eukaryotic cell, *e.g.,* non-treadmilling actin bundles of inner ear stereocilia. Importantly, the correlation between filament stiffness and access to the interstrand space provides a previously unrecognized way of sensing the nucleotide state of actin. It is tempting to speculate that these new functional capacities remain unrecognized but are utilized by host actin-binding proteins.

When the work was ready for submission, a competing Cryo-EM study reported the same association mode of SipA_C_ with actin^45^. While the two works agree on general findings, our study elaborates on many details of these interactions. Indeed, we quantified the competition of SipA_C_ with phalloidin and jasplakinolide and established the correlation between the increased filament stiffness and its susceptibility to depolymerization and severing by cofilin, particularly at stoichiometric doses of SipA_C_. Most importantly, we measured that SipA_C_ binds to F-actin with picomolar K_d_ and established the role of individual domains in this unprecedented binding mechanism.

## Methods

### Protein purification

Skeletal actin was purified from rabbit skeletal muscle acetone powder (Pel-Freeze Biologicals) as previously described^46^. Actin was stored on ice in G-buffer [2 mM Tris-HCl, pH 8.0, 0.2 mM CaCl_2_, 0.2 mM ATP, 0.5 mM β-mercaptoethanol (β-ME), 0.005% sodium azide] for no longer than one month with dialysis after two weeks.

SipA_C_ (a.a. 425-685) was cloned into a pGEX vector with an N-terminal glutathione S-transferase tag (GST) followed by a thrombin cleavage site. Recombinant protein was expressed in BL21(DE3)-pLysS *E. coli* (Agilent Technologies) and purified using glutathione resin (Thermo Fisher). The GST-tag was removed by an on-column cleavage using thrombin (GE Healthcare) at a final concentration of 1 unit/mL. The cleaved SipA_C_ was then passed through a HiPrep 26/60 Sephacryl S-200 HR size-exclusion column (GE Healthcare) using actin polymerization buffer [F-buffer: 10 mM 4-(2-hydroxyethyl)-1-piperazineethanesulfonic acid (HEPES), pH 7.0, 100 mM KCl, 2 mM MgCl_2_, 0.5 mM ethylene glycol-bis(β-aminoethyl ether)-N,N,N′,N′-tetraacetic acid (EGTA), 0.2mM ATP, and 2 mM dithiothreitol (DTT)] that was devoid of ATP. Purified SipA_C_ was flash-frozen in liquid nitrogen and stored at −80 °C.

SipA_C_-ΔArm2 (a.a. 425 – 658) and SipA_C_-Core (a.a. 512 – 658) were cloned with an N-terminal 6xHis-tag into pColdI vector (Takara Bio USA) modified to include a tobacco etch virus (TEV) protease recognition sequence downstream of the 6xHis-tag^47^ using NEBuilder HiFi DNA assembly master mix (New England BioLabs). Sequences were verified by Sanger DNA sequencing [Genomics Shared Resource, The Ohio State University Comprehensive Cancer Center (GSR OSUCCC)]. Recombinant proteins were purified by immobilized metal affinity chromatography (IMAC) as previously described^30^ using HisPur cobalt resin (Thermo Scientific). Before elution, proteins were subjected to on-column cleavage using TEV protease at a 1:20 mole ratio to 6xHis-tagged protein (estimated from resin binding capacity) overnight at 4°C. Purified proteins were dialyzed against ATP-free F-buffer, flash-frozen in liquid nitrogen, and stored at −80°C.

His-tagged ADF was purified as described previously for human cofilins^48^. The 6xHis-tag was removed by TEV protease as described above, and the purified ADF was dialyzed against ADF buffer (10 mM HEPES, pH 7.0, 50 mM KCl, 2 mM MgCl_2_, 0.5 mM EGTA, 2 mM DTT), flash-frozen in liquid nitrogen and stored at −80 °C.

His-tagged mouse CapZ heterodimer construct consisting of α1 and β2 CapZ subunits in pRSFDuet plasmid was a gift from Dr. J. Cooper. 6xHis-tagged CapZ was purified as previously described^30,49^ using HisPur Cobalt resin (Thermo Scientific), and separated from high-molecular weight contaminants by size-exclusion chromatography using a Sephacryl S-200 HR column (Cytiva/GE Healthcare). The purified CapZ was flash-frozen in liquid N_2_ and stored at −80 °C in a buffer containing 20 mM Tris-HCl, pH 8.0, 80 mM KCl, 10 mM DTT, and 20% glycerol.

### Labeling proteins with fluorescent probes

Labeling of actin using Alexa Fluor 488-maleimide and N-(1-pyrene)iodoacetamide (both from Thermo Fisher) was performed as described previously^40^. SipA_C_ constructs were labeled with fluorescein maleimide (FM) as previously described for other proteins^40^.

### Cryo-EM reconstruction

Sample preparation: G-actin was purified from rabbit skeletal muscle. G-actin was stored in G-buffer. Before the experiment, actin was converted to Mg^2+^-ATP G-actin by adding 0.4 mM EGTA and 0.1 mM MgCl_2_. To prepare F-actin/SipA_C_ complex, 5 µM of G-actin was polymerized in G-actin buffer supplemented with 2 mM MgCl_2_ and 50 mM KCl for 24 hours at 4°C. SipA_C_ in 25 mM Tris-HCl, pH 7.9 and 2 mM DTT was added at 5-fold molar excess to actin. 3-µL sample of F-actin/SipA_C_ was applied on discharged lacey carbon grids, and plunge-frozen in liquid ethane using Leica EM GP plunge freezer. Movies were collected on Titan Krios at 300 keV equipped with Gatan K3 direct electron detector. Sampling was 1.08 Å per pixel, total defocus −2.4 to 1.4 µm, with a total dose of around 48 e^-^/Å^2^. Image processing was done in cryoSPARC^50^. Motion- and CTF-corrected images were manually curated for ice thickness and contrast transfer function (CTF) resolution, particles were extracted using 384 box size, followed by a round of 2D classification. To speed up data processing, initial processing was done on a smaller dataset of 169,910 particles (Supplementary Fig. S1). During 3D alignment of particles, a helical symmetry (rise: 27 Å, twist: −167°) was imposed. The first round of 3D classification into 2 classes was done with a bare actin filament and a SipA_C_-bound filament as the initial references. Based on visual inspection of the obtained volumes, a volume made from 90,134 particles showing higher occupancy for SipA_C_ was subjected to 3D variability analysis (3DVA). 1,548,993 particles from a different larger set (384 box size) were then classified into 4 classes based on the volumes obtained from 3DVA. Each class was visually inspected, and a class composed of 349,272 particles was refined using a 112.075 Å rise and −306.344° twist. It was possible to model the actin filament and SipA molecules at 1:2 ratio without steric clashes into the reconstructed volume, supporting the use of these helical parameters. At this step the GSFSC resolution was 5.1 Å. We decided to use all 1,548,993 particles for the reconstruction to improve the resolution. During 3D alignment of 1,548,993 particles, a helical symmetry (rise: 27 Å, twist: −167°) was imposed to get initial alignments. Next, the position and orientation of volumes and particles were adjusted with helical parameters 112.075 Å rise and −306.344° twist and one of the volumes from 3DVA using “Volume alignment tool” in cryoSPARC. Then, non-uniform refinement on 1,548,993 particles was performed, giving a reconstruction of a SipA-decorated actin filament at ∼3.2 Å resolution (Supplementary Fig. S1). Cryo-EM and refinement statistics of the SipAC/F-actin reconstruction are presented in Supplementary Table S3.

SipA crystal structure (PDBID: 1Q5Z) and F-actin cryo-EM model (PDBID: 8D14) were used to build a model of SipA_C_-decorated F-actin using COOT^51^ and ChimeraX^52^. Since the 1Q5Z model contains residues 513-657, the missing residues of the C-terminal tail of SipA_C_ (658-684) were manually built in COOT based on the reconstructed volume. The model was visually inspected and refined in Phenix. Figures were generated using ChimeraX. Mapping of interactions between SipA and F-actin was performed by PBDePISA server^53^. Amino acid residues in SipA model were counted as in the UniProt ID: P0CL52; amino acid sequence for actin model corresponds to UniProt ID: P68135, but in our model the residues were counted using the third aspartic acid as the first residue.

### Analysis of actin’s His73 conformational change upon binding of SipA_C_

Cryo-EM reconstructions of human, rabbit, mouse, chicken, or yeast actin naked or in complex with actin-binding proteins at resolutions better than 3.5 Å were taken for comparison. Models were aligned based on the region between 68^th^ and 93^rd^ amino acid. The list of PDB IDs used for the alignment: 5OOF, 6C1D, 6DJM, 6DJN, 6FHL, 6KLL, 6KLN, 6T1Y, 6T23, 6UPV, 6UPW, 6VAO, 7AHN, 7K20, 7K21, 7P1G, 7PLT, 7PLU, 7PLY, 7PLZ, 7PM3, 7PM5, 7PM6, 7PMD, 7PME, 7PMF, 7PMG, 7PMH, 7PMI, 7PMJ, 7PML, 7R8V, 7R91, 7R94, 7UDT, 8A2R, 8A2S, 8A2T, 8A2U, 8A2Y, 8A2Z, 8D13, 8D14 and 7VVY.

### Co-sedimentation assays

Ca^2+^ in the nucleotide cleft of G-actin was switched to Mg^2+^ by incubating with 0.1 mM MgCl_2_ and 0.5 mM EGTA on ice for 5 min. Actin was polymerized by adding KCl, MgCl_2_, and HEPES (pH 7.0) to final concentrations equal to those of F-buffer, and incubating for 30 min at 25°C. Polymerized F-actin was then mixed to a final concentration of 2.5 μM and titrated with SipA_C_. Reactions were incubated overnight at 4°C, then at 25°C for 1 hour before spinning at 300,000 g for 30 min at 25 °C in a TLA100 rotor in an Optima MAX-TL ultracentrifuge (Beckman Coulter). Supernatants were separated from pellets, resolved on 12% SDS-PAGE, stained with Coomassie Brilliant Blue, and quantified using ImageJ^54^ as described previously^40,47^. The concentration of SipA_C_ in the pellet was quantified by determining the fraction of SipA_C_ pelleted and multiplying it by the known concentration of SipA_C_ in the reaction, allowing for the determination of binding stoichiometry. Data are presented as the average of three technical replicates, error bars show standard deviation (SD) between replicates.

### Equilibrium fluorescence binding assays

To measure the binding affinity of SipA_C_ for F-actin, FM-SipA_C_ was diluted to a final concentration of 1 nM and titrated with F-actin. To limit actin depolymerization prior to the addition of SipA_C_, F-actin was polymerized at 5 µM, and rapidly diluted to the desired concentration in the presence of FM-SipA_C_. For FM-SipA_C_-ΔArm2 and FM-SipA_C_-Core, F-actin was stabilized by incubating with 10 μM phalloidin for 30 min at 25°C. FM-SipA_C_-ΔArm2 and FM-SipA_C_-Core were present at final concentrations of 100 nM and titrated with phalloidin-stabilized F-actin. The final concentration of phalloidin in the reactions was kept at 10 µM to ensure that at least 95% of actin subunits were occupied by phalloidin under all conditions. This step was necessary to prevent depolymerization of F-actin when it was present below its critical concentration. The binding reactions were incubated overnight at 4°C, then at 25°C for 1 hour before measurement. Fluorescence anisotropy (FA) was recorded as previously described^40^ using an Infinite M1000 plate reader (Tecan) with λ_ex_ = 470 nm and λ_em_ = 530 nm. The affinities of TRITC-phalloidin and SiR-actin were measured using 100 nM TRITC-phalloidin (Thermo Fisher) or 10 nM SiR-actin (Cytoskeleton Inc.) titrated with F-actin, and incubated overnight at 4°C followed by 1-hour at 25 °C the next day. Since the fluorescence intensity of TRITC-phalloidin was not changed substantially upon its binding to actin, FA was selected as a reliable reporter of the interaction (Supplementary Fig. S8a,b). In contrast, the fluorescence intensity of SiR-actin increased dramatically upon its interaction with F-actin, making it impossible to accurately measure FA in the free and bound states within the same experiment. (Supplementary Fig. S8c,d). Therefore, fluorescence intensity (FI) was selected as the preferred reporter for SiR-actin binding. For TRITC-phalloidin, FA was recorded at λ_ex_ = 530 nm and λ_em_ = 578 nm, for SiR-actin, FI was recorded at λ_ex_ = 635 nm and λ_em_ = 674 nm. To estimate the binding affinity of the labeled proteins to F-actin, the relative FA or FI changes were fit to the binding isotherm (Equation 1) using Origin 2023 v10.0 (OriginLab Corporation):

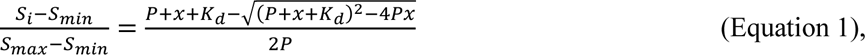

where *S_i_* is the signal (FA for FM/TRITC-labeled components or FI for SiR-actin) for a particular data point, *S_min_* is the minimum signal recorded in that replicate, *S_max_* is the maximum signal recorded in the replicate, *P* is the concentration of labeled binding protein, *x* is the concentration of actin, and *K_d_* is the dissociation constant.

To determine the binding affinity of unlabeled protein for F-actin, we set up a competition assay as described previously^40^. Briefly, labeled components were incubated with F-actin for 30 min, before being incubated with unlabeled proteins of varying concentrations. The mixture was incubated overnight at 4°C, followed by 1 hour at 25°C. FA or FI signals were measured as above. The dissociation constant of the unlabeled protein was estimated by Equation 2:

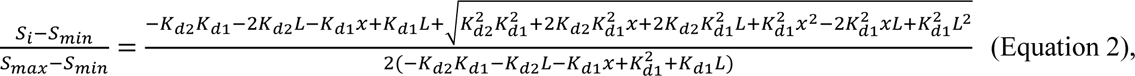

where *K_d1_* is the K_d_ of the labeled component for actin, *K_d2_* is the K_d_ of the unlabeled component for actin, *L* is the actin concentration in the experiment, which is kept constant throughout the titration but is specific to each experiment, and *x* is the concentration of the unlabeled binding partner, which is titrated to the reaction mixture containing the labeled component and actin. The concentrations of actin and labeled protein used in each experiment are given in the Supplementary Table S4. Data are presented as the average of three technical replicates, error bars show SD between replicates.

### F-actin depolymerization assays

Pyrenyl-actin depolymerization assays were performed as described^30^. Depolymerization curves were fit to single exponentials, and the decay rates were plotted against SipA_C_ construct concentrations and fitted with a double exponential equation using Origin 2023 v10.0 (OriginLab Corporation) (Fig. 6e). In addition, the initial decay rates were evaluated by fitting the first 3,000 s of depolymerization curves to a linear equation. The resulting slopes were plotted against SipA_C_ concentration and fitted to a double exponential equation (Fig. 6f). Data are presented as the average of at least three technical replicates, error bars show SD of the mean.

### Cell culture, transient transfections, and microscopy

Human U2OS cells were maintained at 37 °C with 5% CO_2_ in a humidified incubator in Dulbecco’s modified Eagle’s medium (DMEM) supplemented with 10% FBS, L-glutamine, and penicillin-streptomycin. U2OS cells were authenticated by STR profiling using microsatellite genotyping (GSR OSUCCC) and confirmed to be mycoplasma-negative as determined by PCR per published protocol^55^.

SipA constructs were cloned into mammalian expression vectors using NEBuilder assembly (New England Biolabs). SipA full-length (a.a. 1-685), SipA-NT (1-282), SipA-ΔNT (283-685), SipA_C_ (425-685), SipA_C_-ΔArm1 (496-485), SipA_C_-ΔArm2 (425-658), SipA_C_-Core (512-659), and SipA_C_-tandem Core (a.a. 512-664 and a.a.512-659 separated by a linker PAYESGSSASG) were cloned into dCMV-EGFP, a plasmid with a defective CMV promoter optimized for low levels of fluorescent protein expression^56^, as reported previously^32^. Two SipA_C_-Arm2 (a.a. 659-684) constructs were created: 1) in-frame with C-terminal mEmerald-tag cloned into pcDNA3 and 2) with N-terminal TagRFPT cloned into TagRFPT-C1. pmCherry-F-tractin was a gift from Tobias Meyer (Addgene #155218; RRID:Addgene_155218)^57^. pEGFP-F-tractin was a gift from Dyche Mullins (Addgene #58473; RRID:Addgene_58473)^58^.

Cells were co-transfected with F-tractin and various SipA constructs using Lipofectamine 3000 (Thermo Fisher Scientific). Whole-cell Z-stack images of transiently transfected cells were collected using Nikon AX confocal microscope system; maximum intensity projection images were generated using ImageJ software.

### Differential scanning fluorimetry

The effects of phalloidin and SipA_C_ constructs on the thermal stability of F-actin were measured using a CFX Connect Real-Time PCR system (Bio-Rad Laboratories) using Bio-Rad Manager v.3.1 software as previously described^59^. Reactions were carried out in F-buffer supplemented with SYPRO Orange dye (1x final concentration; Invitrogen). 1 µM actin was equilibrated with a range of toxin concentrations and incubated overnight at 4°C before measurement. Melting temperatures (T_m_) were determined by calculating the maximum of the first derivative (dF/dT) of the fluorescence traces. T_m_s of actin were plotted against toxin concentrations using Origin 2023 v.10.0. These data are presented as the average of at least three technical replicates, error bars show SD of the mean.

### F-actin severing assays

F-actin severing by ADF was assessed by TIRF microscopy in chambers made using 22 x 40 mm coverslips and 25 x 75 mm microscope slides (Fisher Scientific), adhered together using strips of double-sided tape. 1 μM Alexa-488-labeled (33%) actin was polymerized in F-buffer containing 50 mM KCl at 25°C for 30 min, then incubated overnight at 4°C to produce aged ADP-enriched actin. SipA_C_ was added to the pre-polymerized actin to a final concentration of 5 or 50 nM and incubated overnight at 4°C. TIRF chambers were blocked by bovine serum albumin (BSA) by 2 consequent washes with 10 μL of high-salt BSA (HS-BSA) solution (1% BSA, 40 mM Tris-HCl, pH 7.5, 480 mM NaCl) followed by two 10-μL washes with low-salt-BSA (LS-BSA) solution (1% BSA, 40 mM Tris-HCl, pH 7.5, 120 mM KCl) and incubated for 5 min^40^. Using cut pipet tips to minimize sheering, filaments were diluted to 0.1 μM in TIRF buffer [10 mM imidazole, pH 7.0, 0.2 mM EGTA, 50 mM KCl, 10 mM ascorbic acid, 2.5 mM protocatechuate-3,4-dioxygenase (PCD), 0.1 μM protocatechuic acid (PCA) purified as described^60^, 100 mM DTT, 0.2 mM ATP, 0.1% BSA] and applied to the chamber. Images were collected with a Nikon Eclipse Ti-E microscope equipped with a TIRF illumination module (with a 15-mW laser), a Nikon CFI Plan Apochromat λ 100× oil objective (NA: 1.45), a perfect focus system (Nikon Instruments), and an iXon Ultra 897 EMCCD camera (Andor Technology). One pixel was equivalent to 160 x 160 nm. After the first 10 frames, 250 nM ADF with 37.5 nM CapZ in TIRF buffer (to prevent reannealing of severed filaments) or CapZ alone as a control were applied to the chamber and images were captured for additional 10 minutes. Using an automated stage, images were captured every 20 s in three separate locations per experiment and in three independent experiments (*i.e.,* nine fields per experimental condition). The time-lapse images were analyzed using ImageJ software by counting severing events of 10 randomly selected filaments in each field of view. The means of cumulative severing events per µm per frame, standard error (SE) and SD of the mean were calculated from nine fields (n = 9) for each experimental condition. Severing was reported as the number of severing events per µm of filament length; error bars represent SE of the mean. For statistical analysis of cumulative severing events per µm of filament length at the endpoint of the experiment (t = 480 s), one-way analysis of variance (ANOVA) followed by Turkey’s post-test was performed using Origin 2023 v10.0 with the *n* value taken as the number of TIRF chamber locations analyzed (*n* = 9) (Supplementary Fig. S5).

### Persistence length analysis

For persistence length measurements, the coverslips and microscope slides were cleaned as described^61^. Coverslips were then dried with nitrogen and layered with 200 µL of 2 mg/mL methoxy-poly (ethylene glycol)-silane (Laysan Bio Inc.) in 80% ethanol. The coverslips were placed in a covered metal dish with their treated sides up and incubated in an oven at 70°C overnight. The coverslips were stored under nitrogen atmosphere in 50 mL conical tubes wrapped in parafilm at −20°C. TIRF chambers were assembled from the treated coverslips and cleaned slides immediately prior to each experiment. To prepare freshly polymerized actin, Ca^2+^ in the nucleotide cleft of Alexa-488-labeled G-actin (3 μM; 33% labeled) was switched to Mg^2+^ by adding 0.1 mM MgCl_2_ and 0.5 mM EGTA and incubating for 2 min. Actin was then polymerized in TIRF buffer either alone or in the presence of saturating concentrations of SipA_C_ (1.5 µM) or SipA_C_-ΔArm2 (3 μM). To evaluate the effects of SipA_C_ on aged, ADP-enriched actin filaments, we prepared ADP-actin filaments as previously described^62,63^. Briefly, Alexa-488-actin (25 μM; 33% labeled) was polymerized for 30 min at 25°C in F-buffer, any remaining ATP was then hydrolyzed with hexokinase (Sigma Aldrich) in the presence of dextrose and Mg^2+^, by incubating for 30 min at 25°C, and continuing incubation on ice overnight to yield ADP-actin. ADP-BeF_3_ actin was prepared by supplementing ADP-actin with 5 mM NaF and 0.1 mM BeSO_4_ and incubating for at least 1 hour at 25°C. For phalloidin-stabilized ADP filaments, Alexa-488-ADP-actin (2 μM total; 33% labeled) was incubated with 2.1 μM phalloidin for 1 hour at 25°C and diluted to a final concentration of 0.1 µM actin immediately prior to the experiment. For SipA_C_-decorated ADP-actin filaments, Alexa-488 ADP-actin filaments (24 μM total; 33% labeled) were incubated with varying concentrations of SipA_C_ for 3 hours at 25°C. The samples were diluted to a final concentration of 0.1 µM actin as above.

Filaments were observed using a Nikon Eclipse Ti-E microscope equipped with a perfect focus system, a TIRF illumination module (with a 15-mW laser), a Nikon CFI Plan Apochromat λ 100× oil objective (NA: 1.45), and a DS-QiMc camera (Nikon). One pixel was equivalent to 65 x 65 nm. Images were captured every 20 s for ∼20 min in three locations in the TIRF chamber per experiment using an automated stage.

Filament contours (“snakes”) were generated using Jfilament^64^ from TIRF time-lapse image files. The resulting snake files were split into individual .dat files. These individual files were then analyzed using the “persistence length analyzer” MatLab script^65^ that determines the average persistence length and SD based on the cumulative curvature distributions of the data. The bootstrapping function was used with an *m* range of 1 to 40, bin size 75 to 110 over 50 iterations. Each analysis used at least 300 .dat files, generated from at least 30 individual actin filaments tracked for 10 consecutive frames. Only filaments no shorter than 10 µm were considered for the analysis.

### Stopped-flow binding and dissociation assays

Time courses of FM-labeled SipA_C_ constructs binding to or dissociating from actin filaments were recorded, and association and dissociation rates were determined by the change in fluorescence anisotropy signal, detected by an SX-20 LED stopped-flow spectrometer (Applied Photophysics) at 25 °C. Excitation of the sample at 470 nm was accomplished by an LED element (Applied Photophysics), and changes in fluorescence anisotropy were measured using two identical 515 nm long-pass colored glass filters (Newport Corporation). For association rates, the reaction was initiated by mixing equal volumes of 100 nM FM-SipA_C_ constructs with equal volumes of F-actin at a range of concentrations in F-buffer. For dissociation rate measurements, an equilibrated mixture of FM-SipA_C_ construct and F-actin was mixed with at least 10-fold excess of unlabeled SipA_C_ construct in F-buffer. The dead-time of the instrument is 1 ms.

Time courses from the association experiments were combined, averaged, and smoothed using Pro-Data SX and Pro-Data viewer (Applied Photophysics) and fit to a single exponential. The resulting *k_obs_* values were plotted against actin concentration in Origin 2023 v10.0 and fit to a linear line, the slope of which yielded the sought *k_on_* values, and the y-intercept was a rough estimate of *k_off_*. All *k_obs_* values are the result of at least 10 association time-course replicates; error bars represent curve fit error. All *k_on_* values are reported with their curve fit errors. *k_off_* values were then measured by observing the dissociation of the fluorescently-labeled constructs from F-actin upon their competition with the excess of unlabeled proteins. The *k_off_* values for SipA_C_-Core and SipA_C_-ΔArm2 were determined by averaging curves from at least ten replicates for two different concentrations of the unlabeled binding partner (at least 60-fold molar excess over the labeled protein and 10-fold molar excess over the concentration of actin). Due to the dual binding mode of SipA_C_, unusually high concentrations of unlabeled SipA_C_ (*i.e.,* 800 and 1000-fold molar excess) were required to effectively displace the labeled protein from F-actin. The dissociation curves of four repetitions at the 1000-fold molar excess of unlabeled SipA_C_ were used (Fig. 4b). The dissociation time courses were averaged, smoothed, fit to a single exponential, and reported together with the curve fit errors derived from the plots produced using the stopped-flow instrument software packages Pro-Data SX and Pro-Data viewer. The experimental *k_off_* values were added to the experimental association graphs to define the intersection of the linear fits of *k_obs_* with the y -axis. The resulting plots maintained the linear dependence on actin concentration and produced the corrected *k_on_* values reported in Fig. 4a.

### Single-molecule TIRFM based dissociation assays

To eliminate the competition effect on the dissociation of SipA_C_ from F-actin, we visualized the spontaneous drop in the number of Alexa-647 labeled SipA_C_ spots associated with Alexa-488 labeled actin filaments in TIRF microscopy (Fig. 4d). TIRF chambers were prepared as described above, except that cleaned and dry coverslips were layered with 200 µL of 2 mg/mL methoxy-poly (ethylene glycol)-silane (Laysan Bio Inc.) and 2 µg/mL Biotin-PEG-Silane (MW 3,400 g/mol, Layson Bio Inc.) in 80% ethanol. SipA_C_ buffer containing 0.01 mg/mL streptavidin and 1% BSA was flowed into the assembled TIRF chamber and incubated for 1 minute, followed by washing with HS-BSA and LS-BSA solutions. To attach F-actin to the substrate and simultaneously stabilize it, SipA_C_ labeled with EZ-Link™ Maleimide-PEG2-Biotin (Thermo Fisher Scientific) was flushed into the chamber at a concentration of 1 µM, and incubated for 5 min, followed by the replacement of the unbound protein with 15 µL of 1x TIRF buffer. Alexa-488 actin was polymerized in the TIRF chamber as described in the persistence length measurement section of the methods. The presence of biotinylated SipA_C_ anchored the Alexa-488 actin filaments to the glass surface and prevented their depolymerization. Alexa-647-labeled SipA_C_ at a concentration of 0.1 nM in TIRF buffer was then added to the TIRF chamber and allowed to decorate actin filaments to the level of individual, clearly distinguishable spots. Unbound Alexa-647-labeled SipA_C_ was washed away by 3 consecutive additions of 15 µL TIRF buffer. The chamber was then sealed with a melted 1:1:1 mixture of Vaseline, lanolin and paraffin (VALAP) to prevent evaporation of the experimental mixture.

To distinguish between SipA_C_ dissociation and photobleaching of the fluorophore (Alexa-647), we monitored drop in fluorescence of Alexa-647-actin (0.5%) copolymerized with Alexa 488-actin (10%) and unlabeled actin (89.5%), stabilized by phalloidin (1.25µM) and biotinylated SipA_C_, which was also used to attach the filaments to the glass surface as described above. Since filaments did not visibly shorten during the experiment duration, we assumed that disappearance of the Alexa-647-actin spots was due to the fluorophore photobleaching. Alexa-647 SipA_C_ dissociation data were corrected for photobleaching and plotted as average of four independent experiments. Error bars are reported as SD of the mean.

### Actin polymerization assays

For polymerization assays, monomeric actin was separated from oligomeric species by passing it through a HiPrep 26/60 Sephacryl S-200 HR size-exclusion column (GE Healthcare) equilibrated in G-buffer. Nuclei-free G-actin from the trailing half of the chromatographic peak was used in the subsequent assays. Actin polymerization was initiated by the addition of 20x EnzChek phosphate assay (Thermo Fisher Scientific) buffer to a final concentration of 1x (50 mM Tris-HCl, pH 7.5, 1 mM MgCl_2_, 0.1 mM NaN_3_) in a 3 mm cuvette (Starna Cells, Inc). The final concentration of actin was 10 µM. When desired, SipA_C_ or SipA_C_-Δarm2 (10 µM final concentration) were added to actin together with the phosphate assay buffer. Polymerization was monitored by changes in light scattering at 90° to the incident light using a PTI QuantaMaster 400 spectrofluorometer (Horiba) at 25 °C. The excitation and emission wavelengths were set to 350 nm with 1.5 nm instrument slit size and ∼2 mm house-made carboard slits installed at the excitation and emission sides of the sample chamber. Prior to the experiment, all solutions were degassed for 1 hour under vacuum. The intensity signal was normalized to the maximum of each condition before averaging, error bars represent SD of the mean. Each curve is the average of at least two independent experiments.

### P_i_ release assays

Inorganic phosphate (P_i_) release from polymerizing F-actin was performed using an EnzChek phosphate assay kit (Thermo Fisher Scientific) according to the manufacturer’s protocol and under conditions identical to those described above for actin polymerization. Polymerization was initiated by adding Sephacryl S-200-purified G-actin to a “pre-reaction” solution in the plate containing the relevant SipA_C_ construct, 2-amino-6-mercapto-7-methyl-purine riboside (MESG) substrate, phosphate assay kit reaction buffer, and purine nucleoside phosphorylase (PNP). After addition of actin, the final volume of the reaction in the plate was 200 μL, with final concentrations of 10 μM actin, 10 μM SipA_C_ construct, 200 μM MESG substrate, 1 unit/mL PNP, 50 mM Tris-HCl, pH 7.5, 1 mM MgCl_2_, 0.1 mM NaN_3_. Reactions were assembled in a 96-well UV-transparent flat-bottom plate (Thermo Fisher Scientific), and reaction progress was monitored by observing absorbance at 360 nm using an Infinite M1000 plate reader (Tecan) with temperature held constant at 25°C. To obtain a strong and reliable signal, it was essential to use 10 μM actin in a 200 μL reaction volume. Measurements were taken every 30 s for 10,000 s. Absorbance traces for actin alone and actin with SipA_C_-Δarm2 showed a decrease in signal after reaching the plateau, which we interpret as spontaneous degradation of the MESG-P_i_ product^66^. For clarity, that portion of the curve was removed. Each curve is the result of at least three technical replicates over at least two independent experiments. Error bars are reported as SD of the mean.

### Comparison of P_i_ release and polymerization rates

Since polymerization curves were shorter than Pi release curves, we extrapolated our data by fitting them to the Hill equation (Equation 3) and subtracted the P_i_ release fits from their respective polymerization counterparts.

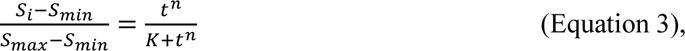

where *t* is the time, *n* is the Hill coefficient, and *K* is a constant. This manipulation produced a plot which shows the fraction of filamentous actin in the ATP or ADP-P_i_ state for any given time point during the experiment, as shown in Fig. 8b.

### Statistical analysis and reproducibility

Graphical data were plotted and analyzed using Origin 2023 v10.0 and Microsoft Excel for Microsoft 365 MSO Version 2307 (Microsoft Corporation). All graph data are presented as the mean ± standard deviation (SD) or standard error (SE) of the mean as indicated in the corresponding method sections. The number of repetitions (n) is indicated in the method section for each technique. For statistical analysis of cumulative severing events of actin filaments, one-way analysis of variance (ANOVA) followed by Turkey’s post-test was performed using Origin 2023 v10.0 with the *n* value taken as the number of TIRF chamber locations analyzed (*n* = 9). For statistical analysis of persistence length measurements, one-way ANOVA followed by Turkey’s post-test was applied using a browser-based calculator (https://statpages.info/anova1sm.html) with the *n* value taken as the number of filaments measured for each experiment (n=30 – 45).

## Author Contributions Statement

E.N. and E.H.E. performed cryo-EM and analysis. L.A.R. and D.S.K. performed biochemistry and analysis. E.K. performed cell culture experiments. All authors wrote and edited the manuscript.

## Acknowledgements

We would like to thank Harper Smith for help with the persistence length analysis, and Kotaro Nakanishi for providing access to the Nikon AX confocal microscope. This work was supported by NIH R35GM122510 (to E.H.E.) and NIH R01GM114666 and R01GM145813 (to D.S.K.). Transmission electron micrographs were recorded at the University of Virginia Molecular Electron Microscopy Core facility (RRID: SCR_019031), which is supported in part by the School of Medicine. In addition, the Titan Krios (SIG S10-RR025067), Falcon II/3EC direct detector (SIG S10-OD018149), and K3/GIF (U24-GM116790) were purchased in part or in full with the designated NIH grants.

## Data availability statement

The cryo-EM map of SipA_C_/F-actin has been deposited with accession code EMD-42161 in the EMDB. The coordinates used for EM analysis have been deposited in the PDB with accession code 8UEE.

