## Supplementary Information for "Stabilization of F-actin by *Salmonella* effector SipA resembles the structural effects of inorganic phosphate and phalloidin"

*Supplementary Figures: S1-S8*

*Supplementary Tables: S1-S5*

*Supplementary Movie Legend: S1*

*Supplementary Figures:*

**Supplementary Figure S1**

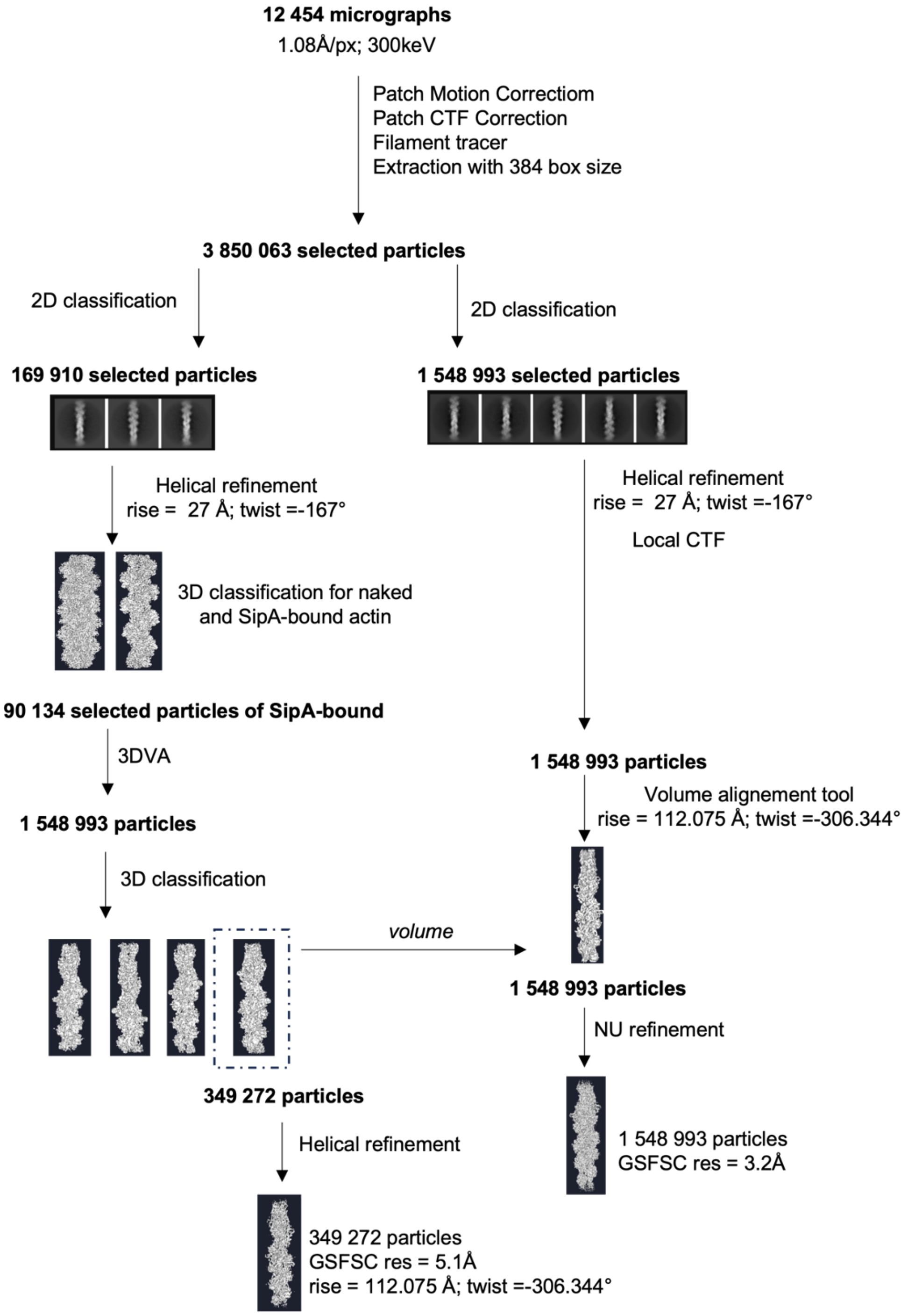


**Supplementary Fig. S1 Cryo-EM data-processing workflow.** Schematic of pre-processing, classification and refinement procedures used to generate the map obtained in this study.

**Supplementary Figure S2**

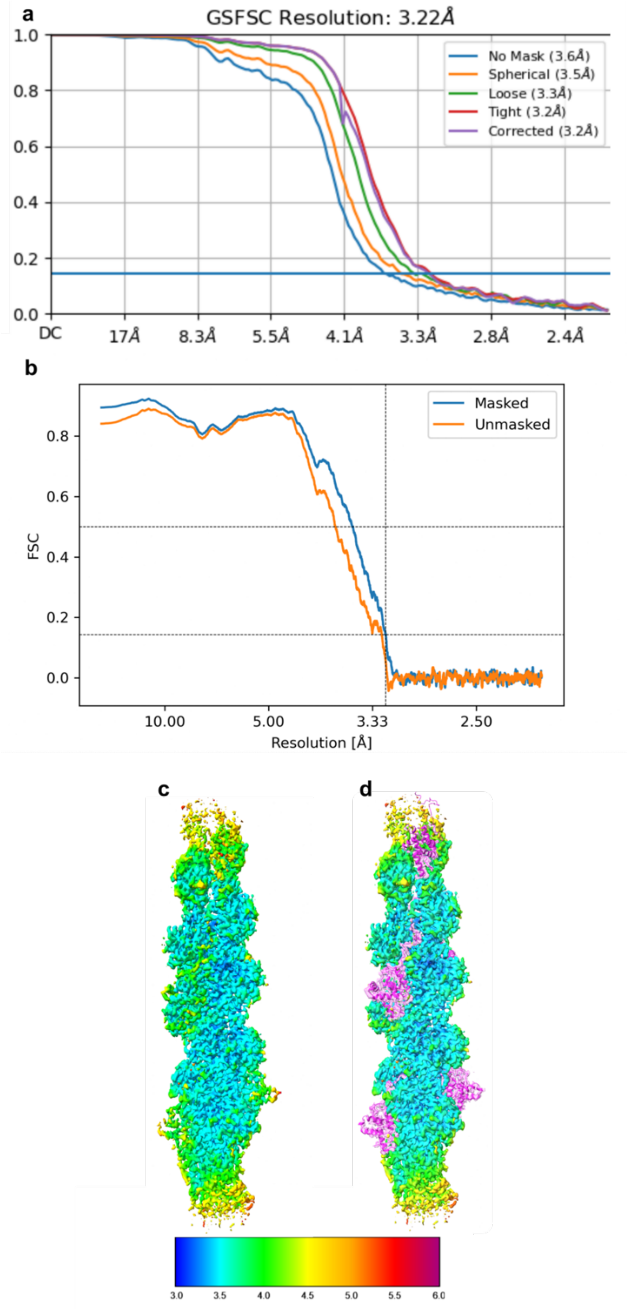


**Supplementary Fig. S2 Fourier Shell Correlation (FSC) calculations, and local resolution estimation of the cryo-EM maps for SipA$_C$/F-actin reconstruction. a,** The map:map FSC calculation (0.143 cutoff). **b,** The model:map FSC calculation (0.5 cutoff). **c,d,** The sharpened cryo-EM density of SipA$_C$/F-actin colored based on local resolution (**c**), and with regions where SipA$_C$ binds (**d**) to show that resolution of reconstruction is the highest for F-actin and decreases towards the outside and at SipA$_C$ binding regions. Values in the color scale are in Å.

**Supplementary Figure S3**

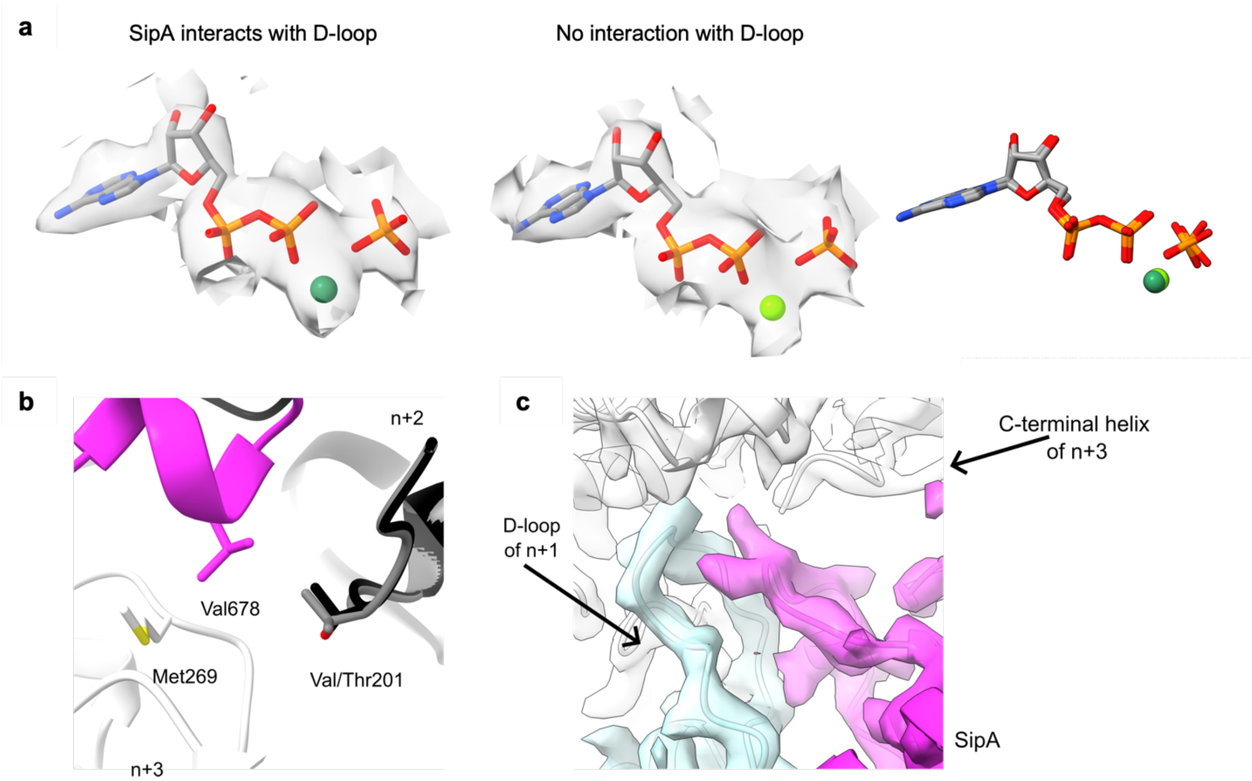


**Supplementary Fig. S3 The effect of $SipA_C$ binding on the occupancy of ADP, $P_i$ and $Mg^{2+}$ in the nucleotide-binding cleft and on the D-loop conformation. a,** Density around ADP, $P_i$ and $Mg^{2+}$ in the nucleotide-binding cleft of the subunits where $SipA_C$ interacts with D-loop (left) and where there is no $SipA_C$ bound to D-loop (middle). There is almost no difference in position of ADP, $Mg^{2+}$ and $P_i$ between these two chains (right). Magnesium ion is colored in green, oxygen atoms are red, phosphate atoms are orange, nitrogen is blue and carbon is grey. **b,** Valine to threonine change in cytoplasmic actin at position 201 does not affect the hydrophobic environment provided by Met269 of actin (in white) and Val678 of SipA (in magenta). The hydroxyl group of threonine can be oriented away from that site, and the methyl groups of valine or threonine can contribute to the formation of a hydrophobic environment. **c,** The D-loop in protomer *n+1* (light blue) adopts a closed state, and the C-terminus of protomer *n+3* (white) adopts a helical conformation when $SipA_C$ (magenta) is bound to actin.

**Supplementary Figure S4**

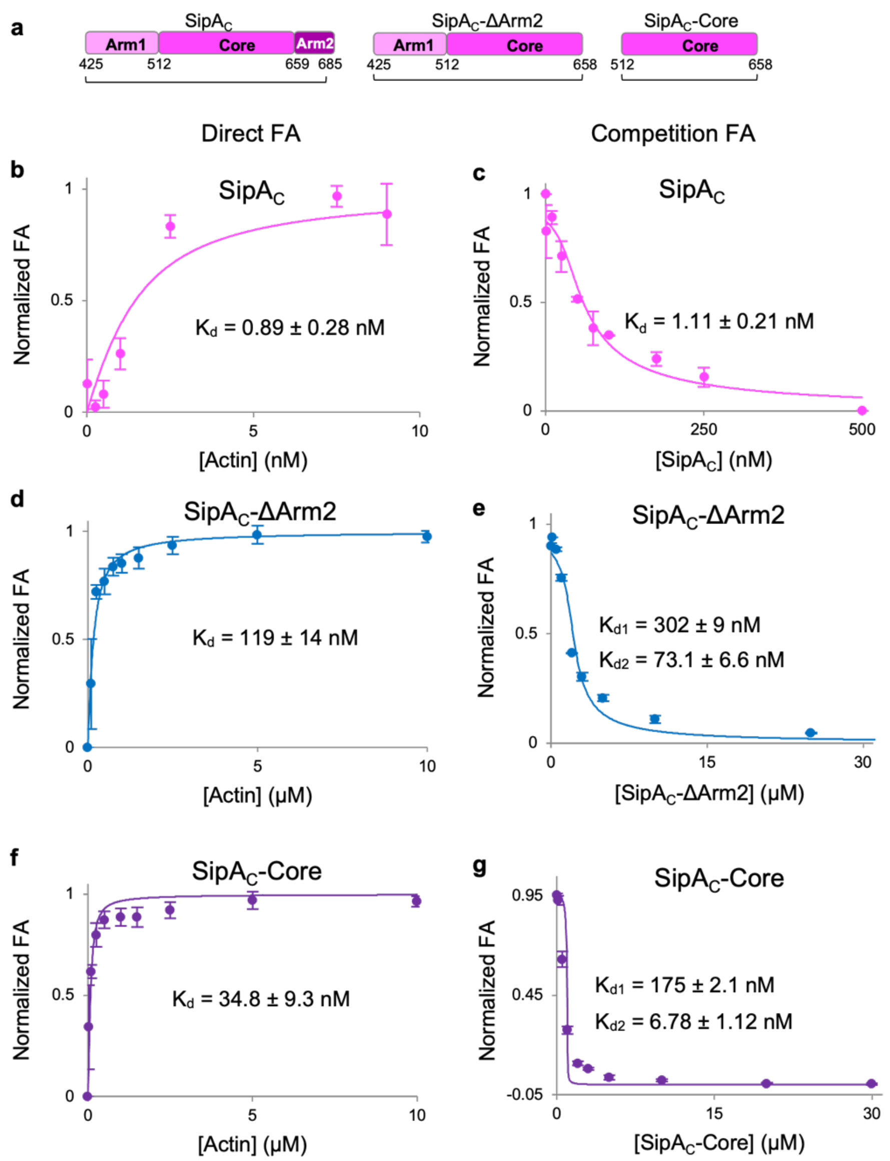


**Supplementary Fig. S4 Arm1 inhibits F-actin binding by SipA$_C$. a,** Domain schematics of SipA$_C$ constructs used for equilibrium FA assays. **b-g,** FA analysis of binding of FM-labeled SipA$_C$ constructs to F-actin (**b, d, f,**) and their displacement by the unlabeled constructs (**c, e, g**). Actin filaments were stabilized by phalloidin in the binding experiments with SipA$_C$-ΔArm2 (**d**) and SipA$_C$-Core (**f**), but not with SipA$_C$, which competes with phalloidin. Phalloidin was not used in the displacement experiments since actin concentration was above critical.

**Supplementary Figure S5**

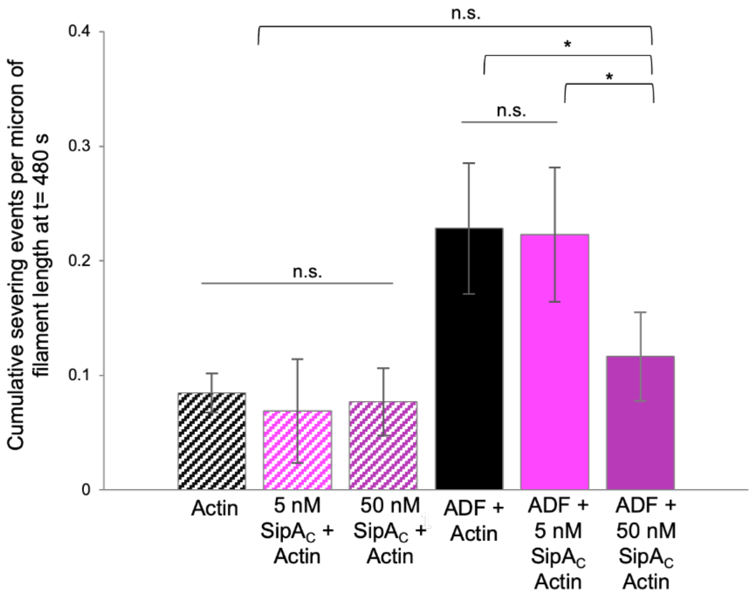


**Supplementary Fig. S5 SipA$_C$ inhibits F-actin severing by ADF.** The average cumulative severing events per µm of filament length at the end-point of the experiment from Fig. 6g,h are plotted with their respective standard deviations. The presence of significant differences between averages were determined by ANOVA analysis (see Supplementary Table S5).

**Supplementary Figure S6**

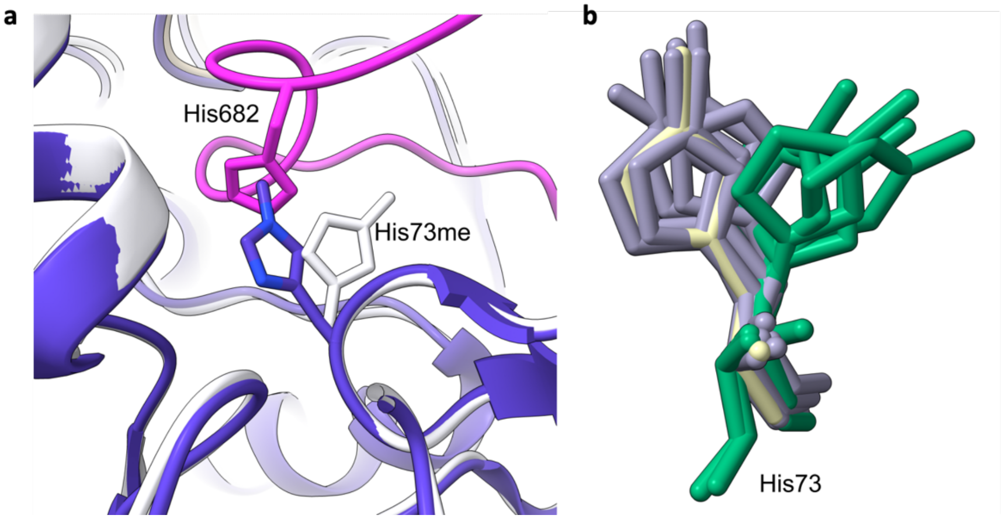


**Supplementary Fig. S6 Binding of Arm2 of SipA$_C$ reorients His73me of actin. a,** His682 on SipA$_C$ (magenta) causes His73me on actin to rotate from its bare actin state (blue) to a new conformation when SipA$_C$ is bound (white). **b,** Conformations of actin His73 when Arm2 is absent (grey) or inserted between actin strands (green). Conformation of His73 from bare actin (PDBID: 6DJN) is shown in yellow.

**Supplementary Figure S7**

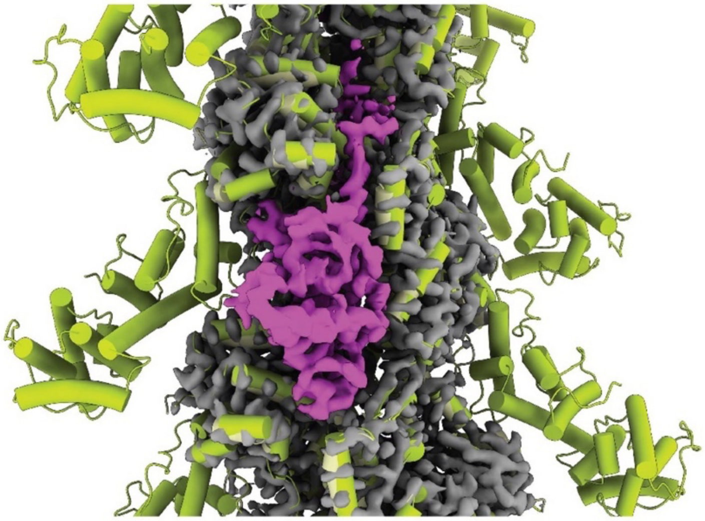

**Supplementary Fig. S7 SipA$_C$ is not in steric conflicts with actin-binding domain of plastin when bound to F-actin.** SipA$_C$/F-actin complex (PDB: 8UEE) aligned to F-actin complex with plastin 2 actin-binding domain 2 (ABD2; PDB:6VEC) shows no steric clash between SipA$_C$ and ABD2. SipA$_C$ is in magenta, actin is in grey, ABD2 of plastin is in yellow.

**Supplementary Figure S8**

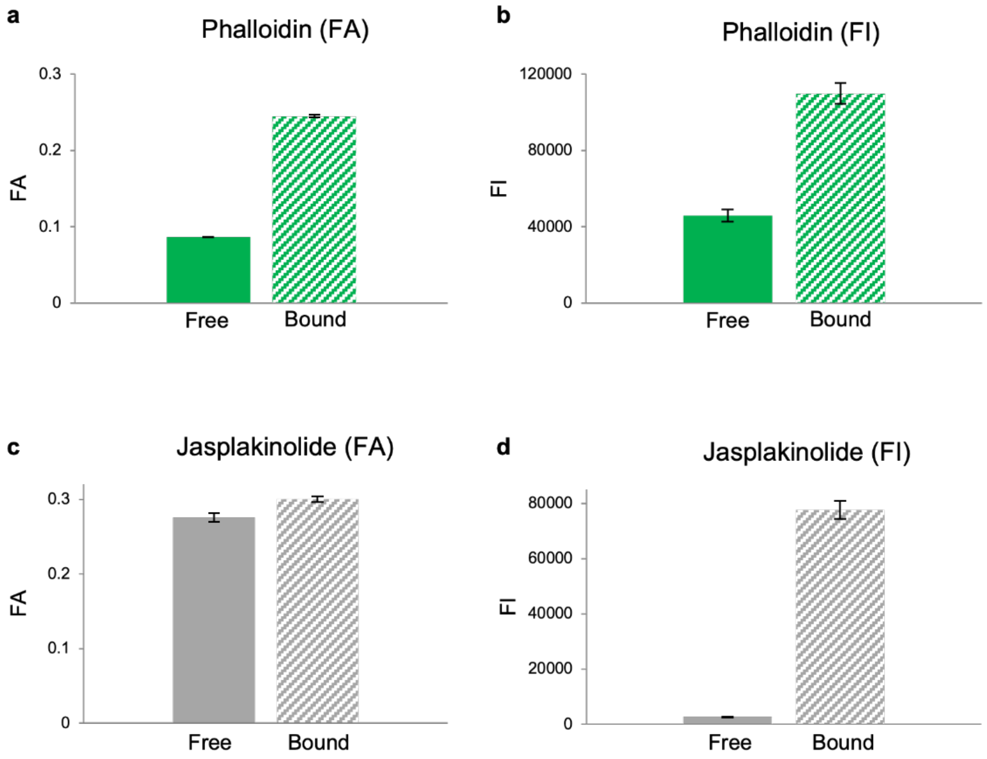


**Supplementary Fig. S8 Comparison of FA and FI as reporters for detecting binding of TRITC-phalloidin and SiR-actin to F-actin. a,b,** Both, the FA (**a**) and FI (**b**) signals of TRITC-phalloidin show an increase in the presence of F-actin. **c,d,** The change in the FA signal (**c**) of SiR-actin (fluorescent derivative of jasplakinolide) upon binding to actin cannot be measured reliably due to a large difference in FI signal (**d**) in the free and bound states. The FI signal was therefore selected as a reliable reporter for SiR-actin binding to actin.

*Supplementary Tables:*

**Supplementary Table S1. Summary of kinetic and equilibrium parameters of SipA$_C$ constructs.** The equilibrium $K_d$ values were derived from competition experiments by allowing the fitting of the $K_d$ values for both labeled and unlabeled components. $K_d$ values of kinetic experiments were calculated using the $k_{on}$ and $k_{off}$ values measured in stopped-flow experiments (for SipA$_C$-ΔArm2 and SipA$_C$-core) and TIRFM assays (for SipA$_C$). The $K_d$ values measured in competition equilibrium and in kinetic experiments reasonably agree, except for SipA$_C$, whose affinity in the equilibrium experiment is underestimated by ~ 9-fold due to the limitations of the approach: *i)* the necessity to use actin at concentrations well below critical and *ii)* insufficient sensitivity of the instrument at sub-nanomolar concentrations preventing lowing SipA$_C$ concentrations for more accurate measurements.

| | $K_d$ (equilibrium) | | | | |
|---|---|---|---|---|---|
| | FM-labeled | unlabeled | $k_{on}$ | $k_{off}$ | $K_d$ (kinetics) |
| SipA$_C$ | 0.89 ± 0.28 nM | 1.11 ± 0.21 nM | 3.8 ± 0.4 $\mu M^{-1}s^{-1}$ | $3.80x10^{-4} \pm 0.9x10^{-4}$ $s^{-1}$ | 100 pM |
| SipA$_C$-ΔArm2 | 302 ± 9 nM | 73.1 ± 6.6 nM | 1.4 ± 0.1 $\mu M^{-1}s^{-1}$ | 0.587 ± 0.003 $s^{-1}$ | 420 nM |
| SipA$_C$-core | 175 ± 21 nM | 6.78 ± 1.12 nM | 3.7 ± 0.2 $\mu M^{-1}s^{-1}$ | 0.391 ± 0.001 $s^{-1}$ | 106 nM |

**Supplementary Table S2. ANOVA analysis of actin filament persistence length measurements.** One-way ANOVA using a Turkey's post-test was performed on data obtained from persistence length measurements of actin filaments in TIRF.

| Experiment 1 | Mean $L_p$ ± SD | Experiment 2 | Mean Lp ± SD | ΔMean | *P* | |
|---|---|---|---|---|---|---|
| Actin + $SipA_C$ | 22.5 ± 1.5 | Actin | 14.6 ± 1.2 | 7.9 | <0.0001 | |
| Actin + $SipA_C$-ΔArm2 | 15.7 ± 1.2 | Actin | 14.6 ± 1.2 | 1.1 | 0.0032 | |
| ADP-Actin | 12.0 ± 1.1 | Actin | 14.6 ± 1.2 | -2.6 | <0.0001 | |
| BeF3-Actin | 13.8 ± 1.4 | Actin | 14.6 ± 1.2 | -0.8 | 0.178 | n.s. |
| ADP-Actin + $SipA_C$ (2:1) | 17.7 ± 1.5 | Actin | 14.6 ± 1.2 | 3.1 | <0.0001 | |
| ADP-Actin + phalloidin (1:1) | 17.8 ± 1.2 | Actin | 14.6 ± 1.2 | 3.2 | <0.0001 | |
| ADP-Actin + $SipA_C$ (20:1) | 13.5 ± 1.2 | Actin | 14.6 ± 1.2 | -1.1 | 0.009 | |
| ADP-Actin + $SipA_C$ (10:1) | 15.5 ± 1.0 | Actin | 14.6 ± 1.2 | 0.9 | 0.0728 | n.s. |
| Actin + $SipA_C$-ΔArm2 | 15.7 ± 1.2 | Actin + $SipA_C$ | 22.5 ± 1.5 | -6.8 | <0.0001 | |
| ADP-Actin | 12.0 ± 1.1 | Actin + $SipA_C$ | 22.5 ± 1.5 | -10.5 | <0.0001 | |
| BeF3-Actin | 13.8 ± 1.4 | Actin + $SipA_C$ | 22.5 ± 1.5 | -8.7 | <0.0001 | |
| ADP-Actin + $SipA_C$ (2:1) | 17.7 ± 1.5 | Actin + $SipA_C$ | 22.5 ± 1.5 | -4.8 | <0.0001 | |
| ADP-Actin + phalloidin (1:1) | 17.8 ± 1.2 | Actin + $SipA_C$ | 22.5 ± 1.5 | -4.7 | <0.0001 | |
| ADP-Actin + $SipA_C$ (20:1) | 13.5 ± 1.2 | Actin + $SipA_C$ | 22.5 ± 1.5 | -9 | <0.0001 | |
| ADP-Actin + $SipA_C$ (10:1) | 15.5 ± 1.0 | Actin + $SipA_C$ | 22.5 ± 1.5 | -7 | <0.0001 | |
| ADP-Actin | 12.0 ± 1.1 | Actin + $SipA_C$-ΔArm2 | 15.7 ± 1.2 | -3.7 | <0.0001 | |
| BeF3-Actin | 13.8 ± 1.4 | Actin + $SipA_C$-ΔArm2 | 15.7 ± 1.2 | -1.9 | <0.0001 | |
| ADP-Actin + $SipA_C$ (2:1) | 17.7 ± 1.5 | Actin + $SipA_C$-ΔArm2 | 15.7 ± 1.2 | 2 | <0.0001 | |
| ADP-Actin + phalloidin (1:1) | 17.8 ± 1.2 | Actin + $SipA_C$-ΔArm2 | 15.7 ± 1.2 | 2.1 | <0.0001 | |
| ADP-Actin + $SipA_C$ (20:1) | 13.5 ± 1.2 | Actin + $SipA_C$-ΔArm2 | 15.7 ± 1.2 | -2.2 | <0.0001 | |
| ADP-Actin + $SipA_C$ (10:1) | 15.5 ± 1.0 | Actin + $SipA_C$-ΔArm2 | 15.7 ± 1.2 | -0.2 | 0.9993 | n.s. |
| $BeF_3$-Actin | 13.8 ± 1.4 | ADP-Actin | 12.0 ± 1.1 | 1.8 | <0.0001 | |
| ADP-Actin + $SipA_C$ (2:1) | 17.7 ± 1.5 | ADP-Actin | 12.0 ± 1.1 | 5.7 | <0.0001 | |
| ADP-Actin + phalloidin (1:1) | 17.8 ± 1.2 | ADP-Actin | 12.0 ± 1.1 | 5.8 | <0.0001 | |
| ADP-Actin + $SipA_C$ (20:1) | 13.5 ± 1.2 | ADP-Actin | 12.0 ± 1.1 | 1.5 | 0.0002 | |
| ADP-Actin + $SipA_C$ (10:1) | 15.5 ± 1.0 | ADP-Actin | 12.0 ± 1.1 | 3.5 | <0.0001 | |
| ADP-Actin + $SipA_C$ (2:1) | 17.7 ± 1.5 | $BeF_3$-Actin | 13.8 ± 1.4 | 3.9 | <0.0001 | |
| ADP-Actin + phalloidin (1:1) | 17.8 ± 1.2 | $BeF_3$-Actin | 13.8 ± 1.4 | 4 | <0.0001 | |
| ADP-Actin + $SipA_C$ (20:1) | 13.5 ± 1.2 | $BeF_3$-Actin | 13.8 ± 1.4 | -0.3 | 0.9925 | n.s. |
| ADP-Actin + $SipA_C$ (10:1) | 15.5 ± 1.0 | $BeF_3$-Actin | 13.8 ± 1.4 | 1.7 | <0.0001 | |
| ADP-Actin + phalloidin (1:1) | 17.8 ± 1.2 | ADP-Actin + $SipA_C$ (2:1) | 17.7 ± 1.5 | 0.1 | 1 | n.s. |
| ADP-Actin + $SipA_C$ (20:1) | 13.5 ± 1.2 | ADP-Actin + $SipA_C$ (2:1) | 17.7 ± 1.5 | -4.2 | <0.0001 | |
| ADP-Actin + $SipA_C$ (10:1) | 15.5 ± 1.0 | ADP-Actin + $SipA_C$ (2:1) | 17.7 ± 1.5 | -2.2 | <0.0001 | |
| ADP-Actin + $SipA_C$ (20:1) | 13.5 ± 1.2 | ADP-Actin + phalloidin (1:1) | 17.8 ± 1.2 | -4.3 | <0.0001 | |
| ADP-Actin + $SipA_C$ (10:1) | 15.5 ± 1.0 | ADP-Actin + phalloidin (1:1) | 17.8 ± 1.2 | -2.3 | <0.0001 | |
| ADP-Actin + $SipA_C$ (10:1) | 15.5 ± 1.0 | ADP-Actin + $SipA_C$ (20:1) | 13.5 ± 1.2 | 2 | <0.0001 | |

**Supplementary Table S3. Cryo-EM and refinement statistics of $SipA_C$/F-actin reconstruction.**

| **Parameter** | **$SipA_C$/F-actin** |
|---|---|
| **Data collection and processing** | |
| Voltage (kV) | 300 |
| Electron exposure ($e^{-}$/Å$^{-2}$) | 48 |
| Pixel size (Å) | 1.08 |
| Particle images (n) | 1 548 993 |
| Shift (pixel) | 32 |
| **Helical symmetry** | |
| Point group | C1 |
| Helical rise (Å) | 112.075 |
| Helical twist (°) | -306.344 |
| Map resolution (Å) | 3.2 |
| Map:map FSC (0.143) | 3.2 |
| Model:map FSC (0.5) | 3.6 |
| Model:map FSC (0.143) | 3.2 |
| d99 | 3.4 |
| **Refinement and model validation** | |
| Ramachandran Favored (%) | 99.10 |
| Ramachandran Outliers (%) | 0.0 |
| RSCC | 0.81 |
| Clashscore | 6.69 |
| Bonds RMSD, length (Å) | 0.004 |
| Bonds RMSD, angles (°) | 0.615 |
| **Deposition ID** | |
| PDB (model) | 8UEE |
| EMDB (map) | EMD-42161 |

**Supplementary Table S4. Summary of experimental conditions used in equilibrium competition assays.**

| Experiment | [Labeled component] (nM) | [Actin] (μM) |
|---|---|---|
| FM-$SipA_C$ competition with unlabeled $SipA_C$ | 1 | 0.039 |
| FM-$SipA_C$-ΔArm2 competition with unlabeled $SipA_C$-ΔArm2 | 100 | 2 |
| FM-$SipA_C$-Core competition with unlabeled $SipA_C$-Core | 100 | 1 |
| TRITC-phalloidin competition with unlabeled $SipA_C$ | 100 | 0.55 |
| SiR-actin competition with unlabeled $SipA_C$ | 10 | 0.25 |
| SiR-actin competition with unlabeled $SipA_C$-Core | 10 | 0.25 |

**Supplementary Table S5. ANOVA analysis of cumulative severing events per µm of actin filament length.** One-way ANOVA using Turkey's post-test was performed on data collected during TIRFM ADF-severing experiments of actin filaments with varying degrees of $SipA_C$ decoration.

| Experiment 1 | Mean cumulative severing per µm ± SD | Experiment 2 | Mean cumulative severing per µm ± SD | ΔMean | *P* | |
|---|---|---|---|---|---|---|
| 5 nM $SipA_C$ + Actin | 0.069 ± 0.045 | Actin | 0.085 ± 0.017 | -0.016 | 0.9784 | n.s. |
| 50 nM $SipA_C$ + Actin | 0.077 ± 0.029 | Actin | 0.085 ± 0.017 | -0.008 | 0.9991 | n.s. |
| 50 nM $SipA_C$ + Actin | 0.077 ± 0.029 | 5 nM SipAC + Actin | 0.069 ± 0.045 | 0.008 | 0.9992 | n.s. |
| ADF + Actin | 0.23 ± 0.06 | Actin | 0.085 ± 0.017 | 0.143 | <0.0001 | |
| ADF + Actin | 0.23 ± 0.06 | 5 nM SipAC + Actin | 0.069 ± 0.045 | 0.159 | <0.0001 | |
| ADF + Actin | 0.23 ± 0.06 | 50 nM SipAC + Actin | 0.077 ± 0.029 | 0.151 | <0.0001 | |
| ADF + 5 nM $SipA_C$ + Actin | 0.22 ± 0.06 | Actin | 0.085 ± 0.017 | 0.138 | <0.0001 | |
| ADF + 5 nM $SipA_C$ + Actin | 0.22 ± 0.06 | 5 nM SipAC + Actin | 0.069 ± 0.045 | 0.154 | <0.0001 | |
| ADF + 5 nM $SipA_C$ + Actin | 0.22 ± 0.06 | 50 nM SipAC + Actin | 0.077 ± 0.029 | 0.146 | <0.0001 | |
| ADF + 5 nM $SipA_C$ + Actin | 0.22 ± 0.06 | ADF + Actin | 0.23 ± 0.06 | -0.006 | 0.9999 | n.s. |
| ADF + 50 nM $SipA_C$ + Actin | 0.12 ± 0.04 | Actin | 0.085 ± 0.017 | 0.032 | 0.6934 | n.s. |
| ADF + 50 nM $SipA_C$ + Actin | 0.12 ± 0.04 | 5 nM SipAC + Actin | 0.069 ± 0.045 | 0.047 | 0.2675 | n.s. |
| ADF + 50 nM $SipA_C$ + Actin | 0.12 ± 0.04 | 50 nM SipAC + Actin | 0.077 ± 0.029 | 0.040 | 0.4639 | n.s. |
| ADF + 50 nM $SipA_C$ + Actin | 0.12 ± 0.04 | ADF + Actin | 0.23 ± 0.06 | -0.112 | <0.0001 | |
| ADF + 50 nM $SipA_C$ + Actin | 0.12 ± 0.04 | ADF + 5 nM SipAC + Actin | 0.22 ± 0.06 | -0.106 | 0.0002 | |

*Supplementary Movie Legend:*

**Supplementary Movie S1. $SipA_C$ protects F-actin against severing by ADF.** F-actin (100 nM of 33%-Alexa-488 labeled F-actin) severing was observed by TIRF microscopy. Images of actin filaments were captured for 8 frames before applying indicated proteins: left panel, 37.5 nM CapZ was added to bare actin; middle panel, 37.5 nM CapZ and 250 nM ADF was added to bare actin; right panel, 37.5 nM CapZ and 250 nM ADF were added to 100 nM F-actin pre-decorated with 5 nM $SipA_C$. CapZ was added to actin filaments in all three experiments to cap barbed ends to prevent reannealing of severed filaments. In the presence of $SipA_C$, very short filaments appear with time that are not the result of severing events within the analyzed frame. They likely represent filaments severed elsewhere in the chamber, which accumulate as their depolymerization is strongly inhibited by $SipA_C$.
